# PASV: Automatic protein partitioning and validation using conserved residues

**DOI:** 10.1101/2021.01.20.427478

**Authors:** Ryan M. Moore, Amelia O. Harrison, Daniel J. Nasko, Jessica Chopyk, Metehan Cebeci, Barbra D. Ferrell, Shawn W. Polson, K. Eric Wommack

## Abstract

**Background:** Increasingly, researchers use protein-coding genes from targeted PCR amplification or direct metagenomic sequencing in community and population ecology. Analysis of protein-coding genes presents different challenges from those encountered in traditional SSU rRNA studies. Most protein-coding sequences are annotated based on homology to other computationally-annotated sequences, which can lead to inaccurate annotations. Therefore, the results of sensitive homology searches must be validated to remove false-positives and assess functionality. Multiple lines of *in silico* evidence can be gathered by examining conserved domains and residues identified through biochemical investigations. However, manually validating sequences in this way can be time consuming and error prone, especially in large environmental studies.

**Results:** An automated pipeline for protein active site validation (PASV) was developed to improve validation and partitioning accuracy for protein-coding sequences, combining multiple sequence alignment with expert domain knowledge. PASV was tested using commonly misannotated proteins: ribonucleotide reductase (RNR), alternative oxidase (AOX), and plastid terminal oxidase (PTOX). PASV partitioned 9,906 putative Class I alpha and Class II RNR sequences from bycatch in a global viral metagenomic investigation with >99% true positive and true negative rates. PASV predicted the class of 2,579 RNR sequences in >98% agreement with manual annotations. PASV correctly partitioned all 336 tested AOX and PTOX sequences.

**Conclusions:** PASV provides an automated and accurate way to address post-homology search validation and partitioning of protein-coding marker genes. Source code is released under the MIT license and is found with documentation and usage examples on GitHub at https://github.com/mooreryan/pasv.

## Background

Next generation DNA sequencing has continued to yield ever larger sequence datasets, enabling researchers to leverage vast amounts of sequence data in addressing a variety of scientific questions from cataloguing variation in human genomes [1] and connecting the gut microbiome with human health [2] to examining the circadian clock in soybean [3] and surveying viruses of the global ocean [4]. For example, sequencing has led to substantial advancements in understanding the community and population biology of microorganisms in nature. Nevertheless, while generation of data continually improves, accurate and comprehensive data analysis remains a challenge for investigations leveraging large sequence datasets.

Building on the example of microbial ecology, for decades researchers have relied on sequence based surveys of stable RNA genes, such as SSU rRNA, as phylogenetic markers for assessing the composition of cellular microbial communities. However, the focus on stable and highly conserved RNA gene sequences for microbial ecology studies has limited researcher’s ability for fine scale delineation of cellular microbial populations from one another [5, 6] and identification of viral populations which do not encode SSU rRNA genes [7]. Use of protein-coding gene sequences as phylogenetic markers for community and population ecology studies can address these shortcomings of SSU rRNA analyses. However, accurate identification of protein-coding genes from either targeted amplicon libraries or shotgun metagenomes remains a significant analytical challenge.

In microbial ecology investigations, both stable rRNA and protein coding marker gene sequences are obtained either through targeted PCR amplification or direct sequencing (i.e., shotgun metagenome sequencing) of environmental DNA. Either approach has limitations that are addressed by the other. Targeted PCR amplification can deeply sample microbial populations within a community, detecting even the rarest of members; however, this approach may miss novel diversity by relying on previously sequenced genes for constructing PCR primers [8, 9, 10]. While every effort is made to ensure marker gene primers capture as much diversity as possible, amplification bias is always present [11]. In contrast, metagenome sequence libraries from shotgun sequencing provide a relatively unbiased picture of microbial diversity, with the caveat of a more limited ability for sampling rare populations [12, 13]. With sequence assembly, this approach also provides the genomic context of marker genes, highly useful information for genome to phenome investigations [14]. Nevertheless, shotgun metagenomics presents significant additional analytical and computational requirements making this approach more expensive and difficult [15, 16]. Furthermore, researchers still must drill down to the level of specific genes within metagenomes, such as those that have undergone extensive biochemical characterization, to uncover interesting biological and ecological patterns from the sequencing data [17, 18, 19, 20]. In the case of either approach, accurately determining the identity of a sequence is critical in preventing subsequent errors in phylogenetic and functional analyses.

Assessing the potential gene functions within a community requires annotation of peptide sequences within metagenomes. Homology-based search tools such as BLAST [21] are the bedrock of sequence annotation, however, functional annotation of proteins based on homology can be error prone [22, 23]. Biochemically annotated proteins are relatively rare in major databases, and usually arise from studies of a few select model organisms [24, 25, 26]. As a result, many environmental sequences are annotated based solely on homology to other computationally annotated environmental sequences rather than to biochemically characterized proteins. Often, such environmental sequence annotations are several steps away from a confident, biochemical annotation, which can quickly lead to inaccuracies resulting from “error percolation” [24].

Furthermore, highly sensitive homology search tools used for annotating and identifying marker genes within metagenomes often have high false positive rates [27]. Identifying false positives in functional annotations is an active area of research and many techniques are available. Machine learning algorithms have been used for identifying false positives based on characteristics of multiple sequence alignments (MSAs) [28, 29]. Active site profiling, or examining the characteristics of regions close to a protein’s active sites, has been used for sensitive and functionally relevant annotations [30, 31, 32, 33].

Even with accurate functional annotations, researchers need a means for predicting if a peptide sequence represents a functional enzyme. While a protein’s function cannot be definitively determined *in silico*, evidence can be gathered by examining active sites, allosteric sites, and other key conserved residues established through biochemical investigations. However, manually validating key residues in thousands of peptide sequences using MSAs is time consuming, especially when considering the large volume of marker gene sequences obtained through amplicon or shotgun metagenome studies [4]. Furthermore, multiple sequence alignment quality degrades as the number of sequences in an alignment increases [34], or when the sequences to be aligned are highly divergent from one another [35].

To address the issue of accuracy in the validation of protein-coding gene sequences, an automated pipeline for protein active site validation (PASV) was developed. PASV provides researchers with a fast and accurate method for validating protein active sites and point mutations in particular genes of interest. Combining multiple sequence alignment with expert domain knowledge in an automated way, PASV more accurately identifies functional protein sequences within large sequence datasets. In this way, PASV can be used as a post-homology search processing step to eliminate most false positive hits and peptides that are likely to be non-functional. Additionally, PASV can be used to partition proteins into groups based on the residues present in functionally important positions of an alignment, such as conserved catalytic residues or residues with interesting biochemical properties (e.g., variants in motif B in DNA polymerase I [17]).

The accuracy of PASV was tested using commonly misannotated proteins: ribonucleotide reductase (RNR), alternative oxidase (AOX), and plastid terminal oxidase (PTOX) [36, 37]. In the first case, PASV was used to identify functional RNRs based on active site residues, and to differentiate Class I alpha and Class II RNRs based on a single amino acid residue. In the second case, PASV was used to distinguish two proteins commonly found in plants, AOX and PTOX, which have been previously shown to be difficult to differentiate with homology search alone, but can be readily partitioned using conserved residues [37].

## Methods

### PASV Pipeline Overview

PASV automates the process of aligning query sequences with a set of reference sequences and subsequently validating key residues and regions within the queries (Fig. 1). PASV is not a homology search tool. Rather it is a post-homology search filtering program. PASV uses a set of user-defined key amino acid residue positions to review alignment columns within multiple sequence alignments (MSAs). Key positions ideally will be residues that are both essential to the protein’s function such as active sites and allosteric binding sites, and highly conserved across the diversity of known protein sequences. In this way, PASV leverages the user’s domain knowledge for automated filtering and validation of functional proteins discovered through homology search. Alternatively, key positions may contain residues that, when mutated, display interesting biochemical properties. PASV automatically bins such amino acid variants, providing information on the functional diversity of a given protein. Finally, PASV can automatically filter out query sequences that fail to span a region of interest (ROI) on the reference sequences.

**Figure 1.**
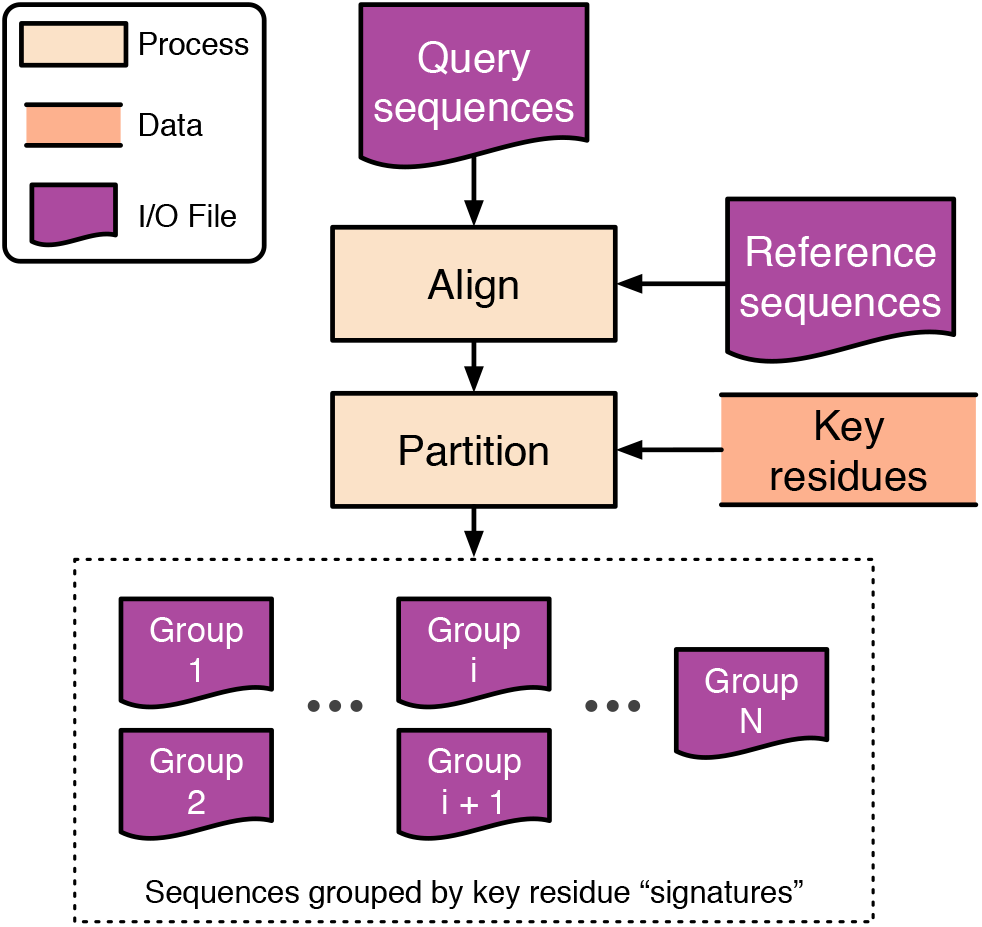
PASV conceptual diagram. PASV individually aligns each query sequence with a user-defined set of reference sequences. Then, columns of the resulting multiple sequence alignment are checked for user-defined key residue positions and, optionally, a region of interest (ROI). Finally, query sequences are partitioned into groups based on the amino acids at each of the key residues and whether the sequence spans the ROI.

Prior to using PASV, users must select a set of reference sequences for the alignment. Special care should be taken when choosing a set of reference sequences, as picking an optimal reference set influences PASV’s accuracy and runtime (see Results and Discussion sections for best practices). Reference sets are tailored to the protein of interest. That is, a set of references chosen for partitioning ribonucleotide reductase (RNR) sequences would not be the same as a set of references used to partition alternative oxidase (AOX) and plastoquinol terminal oxidase (PTOX). In addition to the reference set, which is developed once for a given protein of interest and then reused, the main input to PASV is a set of query protein sequences, generally obtained via a homology search for a protein of interest within a larger sequence dataset. PASV is especially useful in cases where there are many putative protein sequences to validate. For example, using a highly sensitive homology search tool (e.g., BLAST [21], HMMER [38], MMseqs2 [39], or PSI-BLAST [40]) against a metagenome often returns a large set of putative sequences that would be impractical for manual validation. PASV automates sequence validation avoiding time-consuming and potentially error-prone manual validation.

In the PASV pipeline, each query sequence is individually aligned with the reference sequences. PASV abstracts the process of aligning queries with references and identifying residues present in specific columns. Rather than reimplementing MSA algorithms, PASV leverages existing MSA software for aligning queries and reference sequences. It has built-in support for Clustal Omega [41] and MAFFT [42], but other alignment software can be specified at the command line by providing a custom specification.

For each alignment, PASV checks the residues of the query sequence aligning with the user-provided key residue positions in the reference set. The provided key residue positions are interpreted with respect to the original, unaligned first reference sequence. Each query is assigned a key residue “signature” based on these residues. PASV also optionally checks whether each query sequence spans a user-defined region of interest with respect to the reference sequences. Thus, PASV groups query sequences based on the key residue signature, and optionally by ROI spanning status. For example, in the case of RNR, the user may select key residue positions 437, 438, 439, 441, and 462 with respect to the first reference sequence. Then queries will be binned according to the residues that align with the reference sequences at those positions, i.e., their key residue “signatures” (Fig. 2).

**Figure 2.**
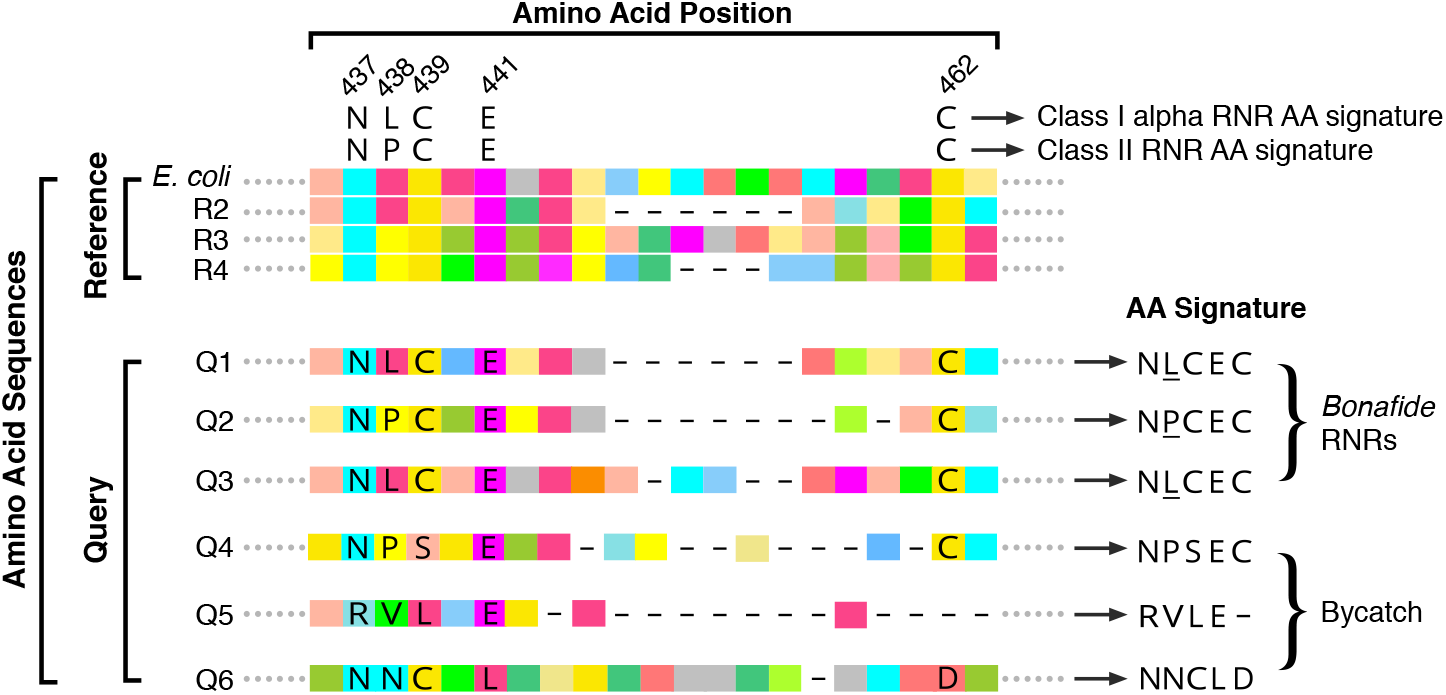
RNR classification and partitioning example. PASV aligns each query sequence individually with all reference sequences (in this case, four references). Labelled positions are the user-specified key residues. The coordinates are specified with respect to the original positions on the unaligned first reference sequence (here, *E. coli*). Each query is assigned a signature based on the residues that align in the same columns as the key residues. In the case of RNR, residues N437, C439, E441, and C462 are required, while residue 438 is diagnostic of RNR class (L438 indicates Class I alpha and P438 indicates Class II). In this example, queries 1, 2, and 3 have NCEC in the correct positions and are considered to be bonafide RNRs. Queries 1 and 3 can be classified as Class I alpha based on L438, whereas query 2 can be classified as Class II based on P438. Queries 4, 5, and 6, do not have the required NCEC signature and are thus considered bycatch.

#### Implementation & source code availability

The PASV pipeline is implemented in Ruby (https://www.ruby-lang.org/), a dynamic, open source programming language. PASV leverages existing multiple sequence alignment software, such as Clustal Omega [41] or MAFFT [42], thus, a multiple sequence alignment program should be installed prior to running PASV. PASV is open-source software (MIT license) and is freely available on GitHub (https://github.com/mooreryan/pasv). Rather than install PASV and its dependencies directly, a Docker image (https://hub.docker.com/r/mooreryan/pasv) and wrapper script (https://github.com/mooreryan/pasv/blob/master/bin/pasv_docker) are also available. PASV v1.3.0 (https://github.com/mooreryan/pasv/releases/tag/v1.3.0) was used for all experiments.

#### PASV result network diagrams

Resulting PASV output files were converted to a node-link network diagram with a custom script (available on the PASV GitHub page) and visualized with Cytoscape v3.7.1 [43].

### Collecting RNR sequences

#### Retrieving RNR sequences from the RNRdb

All available Class I alpha and Class II RNRs were retrieved from the RNRdb on August 20, 2018 [36]. These 66,209 RNR peptide sequences were dereplicated (exact and substring matches) using CD-HIT v4.6 [44], yielding 29,401 representative sequences. Sequences were then divided into closely related groups (clades) as defined by the RNRdb for manual assessment of active site residues and intein removal [23]. From the 29,401 representative sequences, 286 sequences were removed as they lacked one or more of the four residues essential for RNR function (N437, C439, E441, C462 with respect to *Escherichia coli* K12 W3110 ribonucleoside diphosphate reductase 1 alpha subunit, accession no. WP_001075164.1) [45, 46, 47, 48]. The 29,133 remaining RNRs were retained for downstream analysis.

#### RNRdb sequence tree & phylogenetic clustering

To reduce the number of sequences used for building a phylogenetic tree of known RNR peptides, the 29,133 bonafide RNRdb sequences were clustered with MMseqs2 (version e1a1c1226ef22ac3d0da8e8f71adb8fd2388a249) [39] at 75% identity over 80% of the alignment length, resulting in a set of 2,579 peptide clusters. Cluster centroids were aligned with MAFFT v7.427 using the FFT-NS-2 method [42]. Columns of the resulting multiple sequence alignment containing >95% gaps were removed. Finally, FastTree v2.1.10 with double precision arithmetic [50] was used to build the tree, and the resulting tree was midpoint-rooted with a custom Python script (https://github.com/mooreryan/midpoint-root) using ETE Toolkit v3 [51]. Different numbers of phylogenetic RNR clusters were generated by collapsing branches whose lengths were below a threshold using iTOL [49]. Six different clustering scenarios were used representing six levels of phylogenetic granularity (4 clusters: collapsed branch length (BRL) < 3.75; 8 clusters: BRL < 3.1; 14 clusters: BRL < 2.85; 19 clusters: BRL < 2.65; 24 clusters: 2.485; and 29 clusters: BRL < 2.34) (Fig. 3).

**Figure 3.**
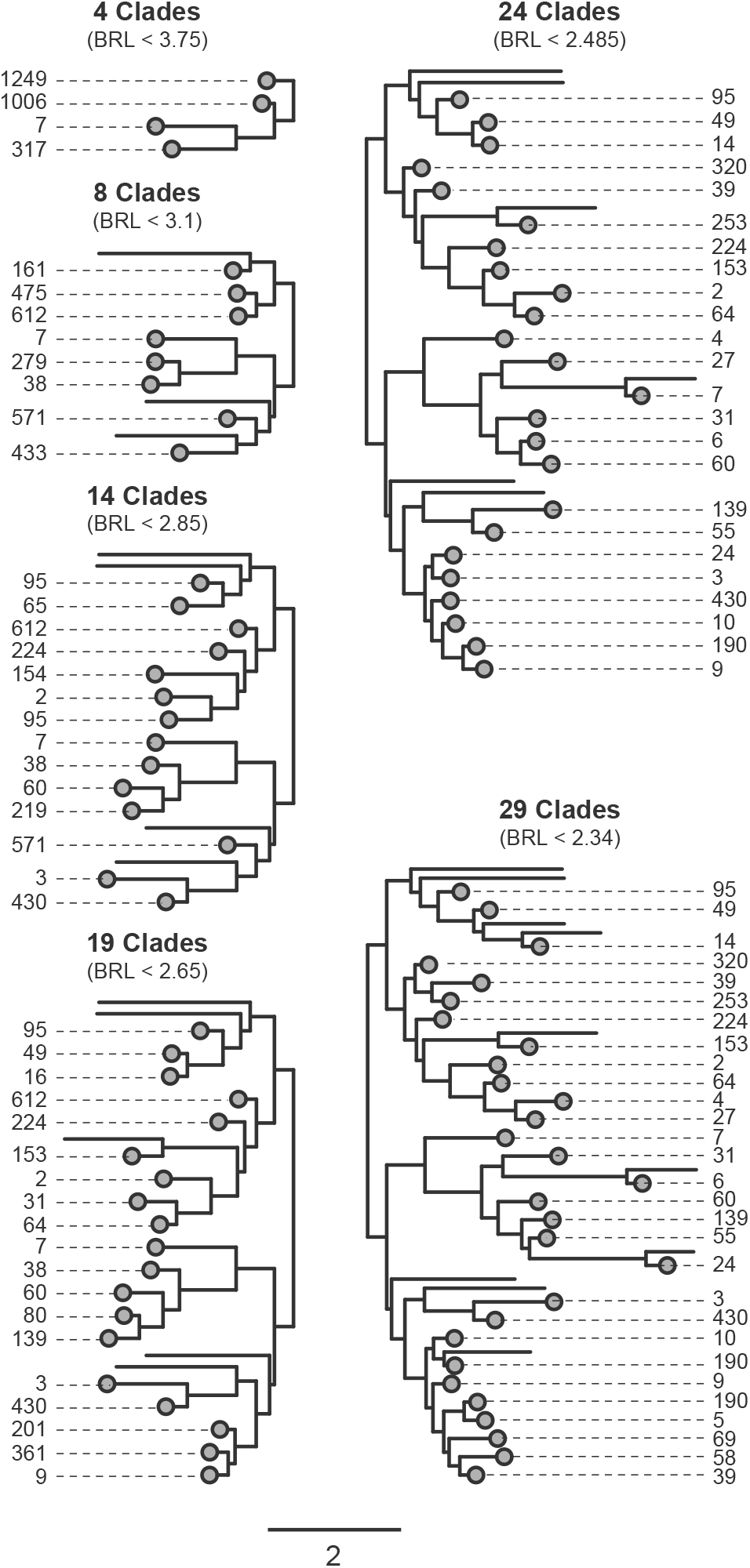
Phylogenetic clustering of ribonucleotide reductase proteins. Ribonucleotide reductases (RNRs) from the RNRdb [36] were clustered with MMseqs2 [39] at 75% identity over 80% of the alignment length. Phylogenetic clusters (grey circles) were created in iTOL [49] by collapsing clades with branch lengths (BRL) less than the amount shown. Leaf labels show the number of sequences within the clade. Branches without grey dots represent singleton clusters, and were not included in the pool of potential reference sequences. Scale bar represents amino acid substitutions per site.

#### Retrieving RNR sequences from the Global Ocean Viromes dataset

The 1,995,784 Global Oceans Virome (GOV) [52] peptides (downloaded from: https://datacommons.cyverse.org/browse/iplant/home/shared/iVirus/GOV/Contigs_set, file last modified 2017-04-23) were searched against RNRdb sequences with MMseqs2 (sensitivity: 7, max-seqs: 1000, num-iterations: 3, start-sens: 1, sens-steps: 7, default e-value cutoff: 0.001, defaults for all other options). This search yielded 12,412 virome sequences. Sequences having fewer than 100 amino acids were removed, leaving 9,906 sequences. These sequences were manually curated using a combination of conserved residues, domains, and phylogenetic placement (as in [23]) resulting in 2,916 bonafide RNRs and 6,990 non-RNRs.

### Reference sets and PASV accuracy

#### Full reference set test

Given that PASV uses MSA for validating key residues, PASV’s accuracy is dependent on the chosen reference set and aligner. An experiment testing 1,920 combinations of reference sets, query sets, and aligners was used to determine those variables most affecting accuracy (Fig. 4). First, randomly selected reference sequence sets were compared to sets where selection was guided by a phylogenetic tree. For phylogenetically selected references, a tree containing 2,579 RNR sequences was partitioned at six levels of granularity (4, 8, 14, 19, 24, and 29 clusters (Fig. 3)). Two approaches were then taken for phylogenetic reference selection. First, phylogenetic reference sets were generated by selecting a single reference sequence from each tree clade (clades defined by various minimum branch lengths (BRL, Fig. 3) to test whether increasing the evenness of representation among rarer or divergent clades would improve PASV accuracy. Second, phylogenetic reference sets were generated by weighting the selection of sequences according to the number of sequences within a cluster (one reference sequence for every 200 sequences in the cluster) (Fig. 3). For each of the phylogenetically selected reference sets (including weighted and unweighted at all six levels of granularity), size-matched, randomly selected reference sets were included as controls. Finally, for each reference set selection criteria (phylogenetic or random, single or multi, reference set size), ten replicates were generated. Each reference set was tested with two aligners, MAFFT v7.427 [42] and Clustal Omega v1.2.4 [41], and two different query sets (RNRdb queries: 100 bonafide RNRs and 100 invalid RNRs missing key functional residues; Global Ocean Virome (GOV) queries: 200 bonafide RNRs and 100 invalid RNRs missing key functional residues). All experiments were run on an Intel(R) Xeon(R) CPU E5-2695 v4 @ 2.10GHz server with 36 cores (2 threads per core) and 512 GB of ram, with PASV set to use 68 threads (i.e., process 68 queries concurrently).

**Figure 4.**
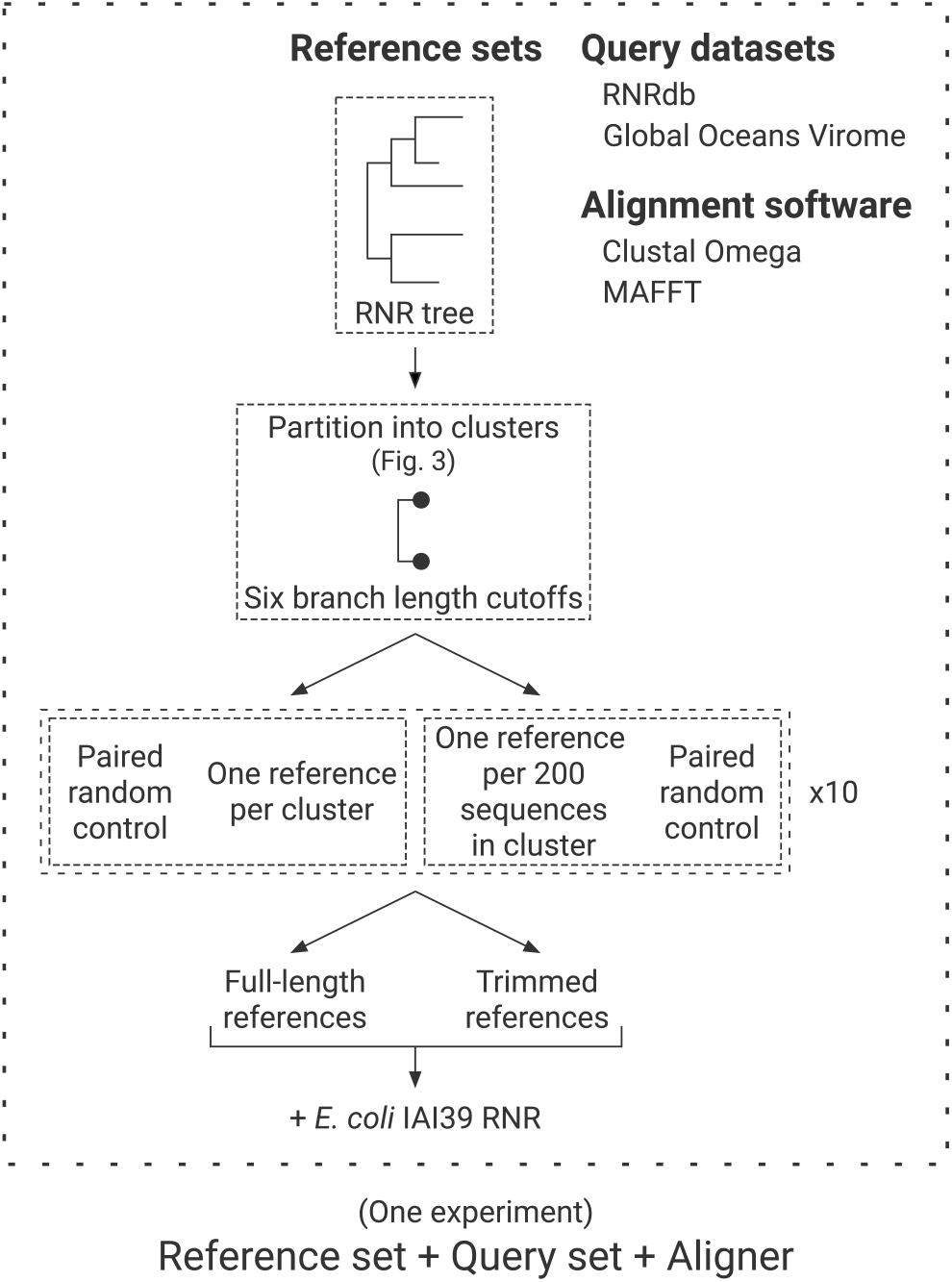
PASV reference set test. Conceptual diagram of the validation experiment testing the effects of reference set, query set, and aligner on PASV accuracy. One experiment is a PASV run with a unique combination of a reference set, a query set, and an aligner. The reference sequence selection strategy (phylogenetically-guided or random), the size of the reference set (numbers of sequences and their distribution across the known diversity of a protein), and the length of reference sequences (full length or smaller region of interest) were tested for their impact on PASV accuracy in correctly identifying manually curated sequences. For each reference set category, 10 random samples (i.e., replicates) were generated. For each reference set, two aligners (Clustal Omega [41], and MAFFT [42]), and two query sets (RNRdb [36] and Global Ocean Virome (GOV) [52]) were run.

In summary, a total of 1,920 experiments were conducted. Six levels of phylogenetic tree partitioning were used (6 reference sets) (Fig. 3), each generating either a single (unweighted) or multiple (weighted) reference sequences per clade (6 * 2 = 12 reference sets). For each of these twelve groups, ten replicates were made (12 * 10 = 120 reference sets). For each of these 120 reference sets, size-matched, randomly selected reference sets were used as controls (120 * 2 = 240 reference sets). For each of these 240 reference sets, both full-length reference sequences, and reference sequences trimmed to the shorter region of interest (ROI, positions 437 - 605, *E. coli* numbering) were tested (240 * 2 = 480 reference sets). For each of these 480 reference sets, two aligners (Clustal Omega or MAFFT) were tested (480 * 2 = 960 reference sets + aligners). For each of these 960 reference set plus aligner pairs, two different query sets (RNRdb or GOV) were tested (960 * 2 = 1,920 experiments) (Fig. 4).

#### Putative GOV RNR queries test

GOV RNR sequences (9,906 sequences) were used to test PASV on a dataset more reflective of an actual use case. Because most of the variables tested in the full reference set test had little effect on PASV accuracy (see Results), and due to the size of the query set, a reduced set of variables was used to generate reference sets. References from three clustering levels (8, 19, 29) with both phylogenetic and random sequence picking were generated in triplicate, yielding 18 reference sets. For the other variables included in the full reference set test, only the top performing options were used in this experiment: Clustal Omega rather than MAFFT, full-length references rather than trimmed, and one sequence per clade vs one sequence for every 200 sequences per clade. All experiments were run on the same server as the full reference set test with PASV set to run 68 concurrent alignment jobs.

#### Data analysis

Data analysis was performed in R v3.6.3 [53] with tidyverse v1.3.0 [54] and ggplot2 v3.3.0 [55]. All true positive and true negative rate linear models were calculated with the lm function in R. Model coefficients were considered significant if their *p*-values were less than 0.05 as reported by the R function summary.lm. All box and whisker plots were made using the geom_boxplot function from ggplot2. All scatter plot regression lines were made using the geom_smooth function from ggplot2 using locally estimated scatterplot smoothing (LOESS, default parameters) with 95% confidence intervals, except for Additional Files 1 and 4 which use linear regression with 95% confidence intervals calculated with geom_smooth using lm. All point jittering was done using the geom_jitterdodge function from ggplot2.

### Analyzing putative and bonafide GOV RNRs

#### GOV RNR trees

The 9,906 putative RNR sequences identified through homology search alone, and the 2,914 PASV-predicted bonafide RNR sequences (using the reference set chosen from the best practices according to the full reference set test and the GOV RNR queries test) were aligned with MAFFT v7.427 FFT-NS-2 [42]. Columns with >95% gaps were removed and a phylogenetic tree was inferred with FastTree v2.1.10 double precision arithmetic [50]. The resulting Newick tree files were visualized with Iroki [56].

#### Annotating GOV tree sequences

Sequences were manually selected from clades containing only non-RNRs (according to manual curation) from the phylogenetic tree containing all 9,906 putative RNRs from GOV. Sequences were searched against National Center for Biotechnology Information’s Conserved Domain Database (NCBI CDD) v3.18 and the top domain hit by e-value was recorded [57]. All sequences that had a mismatch between manual curation and PASV prediction in any of the 18 full GOV experiments were also searched against the conserved domain database using Batch CD-Search [58] and the top domain hit was recorded. In the case that multiple domains were identified, the top hit was recorded for each domain (Additional File 10).

### Partitioning RNR classes

To test PASV’s ability to partition Class I RNR alpha subunit sequences from Class II RNR sequences, the 2,579 clusters from the RNRdb tree (Fig. 3) were used as PASV query sequences with the "best practices" RNR reference set. In addition to the same N437, C439, E441, and C462 key residues (*E. coli* numbering) used in previous experiments, residue L/P438 was also included. Any sequence PASV identified as having N**L**CEC was labeled as a Class I alpha RNR, whereas any sequence with N**P**CEC was classified as a Class II RNR. Any sequences with key residue signatures other than the NLCEC for Class I alpha and NPCEC for Class II were grouped into the “Other” category. The PASV predictions were compared with RNRdb assigned class annotations.

### Partitioning AOX and PTOX

Alternative oxidase (AOX) and plastid terminal oxidase (PTOX) peptide sequences were collected from a recent study [37]. Sequences from supplemental data sheet 1, containing 14 full-length PTOX proteins that were previously erroneously annotated as AOX, and sequences from supplemental data sheet 2, representing trimmed AOX and PTOX sequences, were obtained. Some of the trimmed sequences in supplemental data sheet 2 had accession numbers with which the corresponding full length sequences could be recovered from NCBI databases using the Entrez Direct efetch [59]. Forty-eight full-length AOX and eight PTOX sequences were recovered in this manner. Recovered full length sequences were combined with trimmed sequences yielding a set of 336 query sequences for PASV testing.

The ability of PASV to classify both AOX and PTOX sequences within a mixed set of peptide sequences was tested with two separate PASV runs: once with an AOX reference set (UniProt entry IDs O22048, O22049, and E1CIY3; sequences selected from those manually annotated as AOX in [37]) and once with a PTOX reference set (UniProt entry IDs A0A061GHF5, B9RXE2, and Q56X52; sequences selected from those manually annotated as PTOX in [37]) sequences. In the AOX run, all query sequences were checked for conserved residues from AOX motifs 1 (E233, R234, M235, H236, L237, M238, T239) and 2 (L283, E284, E285, E286, A287), and sequences containing the correct residues were labeled as AOX, while sequences with other residues at these positions were labeled as non-AOX (numbering with respect to sequence O22048) [37]. For the PTOX run, all queries were checked for conserved residues from PTOX motifs 1 (G157, W158, R160, R161) and 2 (H177, H178, L179, L180, M182, E183), and any sequences containing the correct residues were labeled as being PTOX, while sequences with other residues at these positions were labeled as non-PTOX (numbering with respect to sequence A0A061GHF5) [37]. Finally, the sequence labels from the AOX and the PTOX run were combined for the final classification. Two positions were excluded from the motifs that were presented in [37] (159 in motif 1, and 181 in motif 2) as these positions were more variable than the other motif positions.

## Results

### What factors influence PASV accuracy

True positive and true negative rates for PASV validated RNR peptide sequences were explored with linear models. For GOV query sequences, aligner and reference trimming had a significant (*p*-value < 0.05) association with both true positive and true negative rates (Fig. 5). Clustal Omega was associated with an 11.1% increase in true positive rate and a 0.2% decrease in true negative rate as compared to MAFFT. Full length references had a 12.6% increase in true positive rate and a 0.07% increase in true negative rate as compared to references trimmed to the region of interest. While statistically significant according to the linear model, variables associated with true negative rate had negligible effect in practice for GOV queries. For RNRdb queries, when all variables were included as predictors, aligner and reference trimming were both significant predictors of true positive rate. Clustal Omega was associated with a 1.5% increase, and full length references were associated with 2.7% increase in true positive rate. For true negative rate, all variables other than replicate were significant; however, all effects were quite small (< 1.3%).

**Figure 5.**
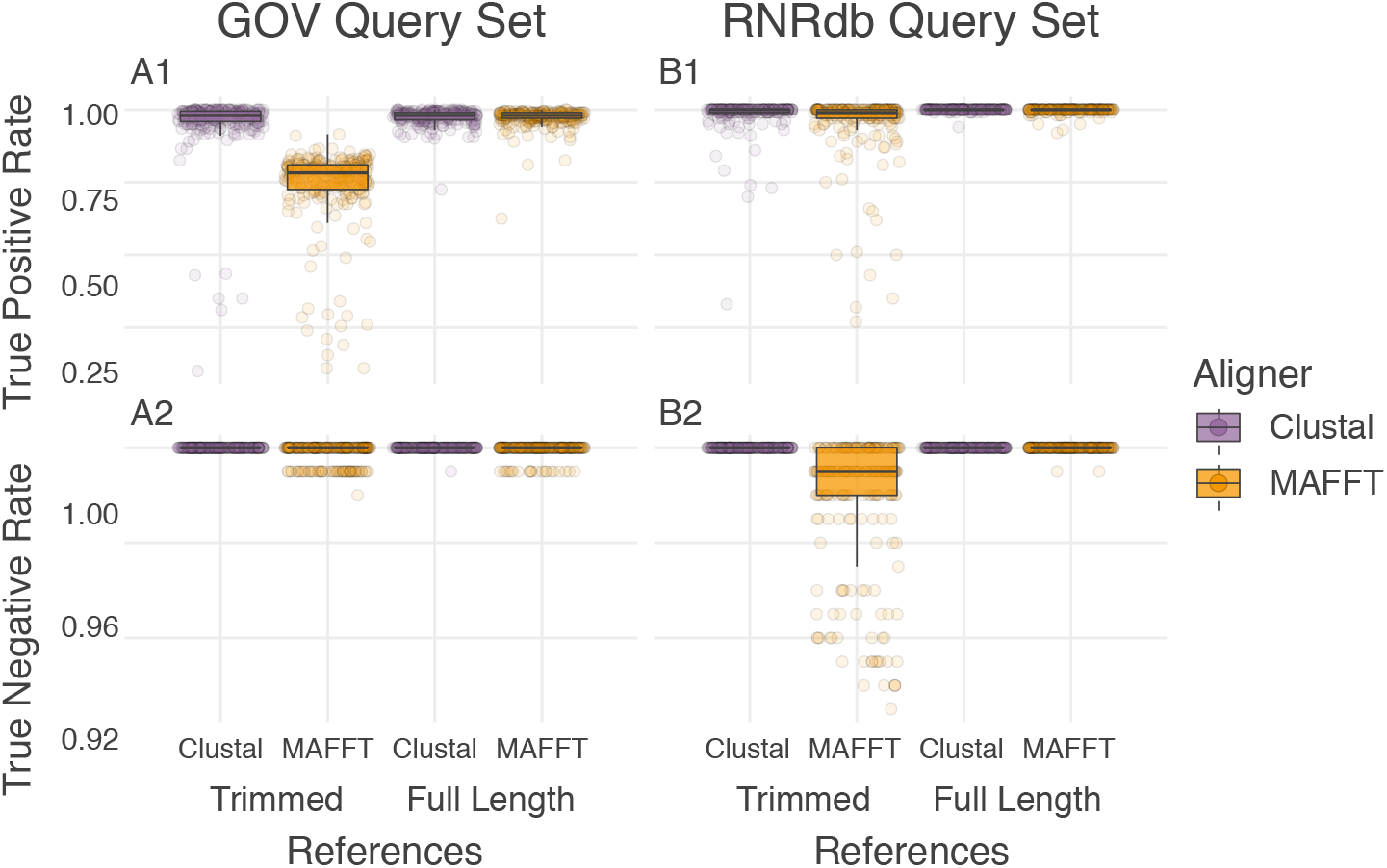
PASV accuracy is influenced by aligner and reference trimming. PASV true positive (A1 & B1) and true negative rates (A2 & B2) across reference sets of RNR peptide sequences. Results are shown for the Global Ocean Virome (GOV) query set (A) and the RNRdb query set (B). Each dot represents a single PASV run (i.e., one reference set with an aligner). Box (showing median and interquartile range (IQR)) and whisker (1.5 x IQR) plots are overlaid. Within each panel, PASV tests are partitioned by reference sequence length (full length references vs. those trimmed to the region of interest) and by multiple sequence aligner (Clustal Omega – purple vs. MAFFT – orange).

Given that full-length references were superior to those trimmed to a ROI (Fig. 5), only full-length references were included in subsequent analysis of covariate effects on PASV accuracy. Full-length reference sets split into groups based on query set (GOV vs RNRdb) and aligner (Clustal Omega vs. MAFFT) were re-run through linear models on the following five remaining covariates: (1) number of tree clusters; (2) number of reference sequences; (3) single or multiple reference sequences chosen per clade (single/multi); (4) random or phylogenetically-guided reference sequence choice (random/phylo); and (5) reference set replicate (Table 1).

**Table 1.** Linear model coefficients with *p*-value < 0.1 for PASV reference set test (full-length references only).

| Model | Variable | Coefficient ± Standard Error | | | |
| --- | --- | --- | --- | --- | --- |
|  |  | GOV-MAFFT | GOV-Clustal | RNRdb-MAFFT | RNRdb-Clustal |
| True positive rate | Intercept | 86.50 ± 1.86 | 97.60 ± 1.69 | 97.30 ± 0.55 | 99.29 ± 0.28 |
| | No. tree clusters<sup>a</sup> | -0.56 ± 0.13 | | -0.12 ± 0.39 | -0.03 ± 0.02 |
| | No. references<sup>b</sup> | 0.78 ± 0.15 | | 0.17 ± 0.45 | 0.05 ± 0.02 |
| | Single vs. multi<sup>c</sup> | 4.77 ± 1.33 | | 1.10 ± 0.39 | |
| | Random vs. phylo<sup>d</sup> | 0.65 ± 0.36 | | 0.24 ± 0.11 | |
|  | Replicate |  |  |  |  |
| True negative rate | Intercept | 99.92 ± 0.19 | 99.99 ± 0.43 | 99.87 ± 0.06 | 100.00 ± 0.00 |
| | No. tree clusters | | | | 0.00 ± 0.00 |
| | No. references | | | 0.01 ± 0.00 | 0.00 ± 0.00 |
|  | Single vs. multi |  |  |  |  |
| | Random vs. phylo | -0.12 ± 0.04 | | | |
| | Replicate | -0.01 ± 0.01 | | | |

<sup>a</sup>Number of tree clusters<sup>b</sup>Number of reference sequences<sup>c</sup>Single or multiple reference sequences chosen per clade<sup>d</sup>Random or phylogenetically-guided reference sequence selection**Table 2.** Confusion matrix of PASV results for 18 references sets against putative GOV RNR sequences.

The PASV true positive rate decreased with the number of tree clusters used in the phylogenetically-guided reference sequence choice approach (Fig. 3), but increased with respect to the number of references for GOV-MAFFT, RNRdb-MAFFT, and RNRdb-Clustal groups (Table 1). When using MAFFT, picking a single reference from each clade as opposed to weighting the number of references by number of sequences in the clade was associated with a significantly higher true positive rate for both GOV and RNRdb query sets; however, this trend was not seen when using Clustal (Table 1). Overall, choosing references randomly (when using MAFFT, but not Clustal) and including more sequences in the reference set were associated with better PASV accuracy. However, the positive effect of the number of reference sequences on true positive rate plateaued after ca. 20 reference sequences (Fig. 6). Additionally, the effect of increasing the number of references is more pronounced with the MAFFT aligner than with Clustal Omega (Fig. 6).

**Figure 6.**
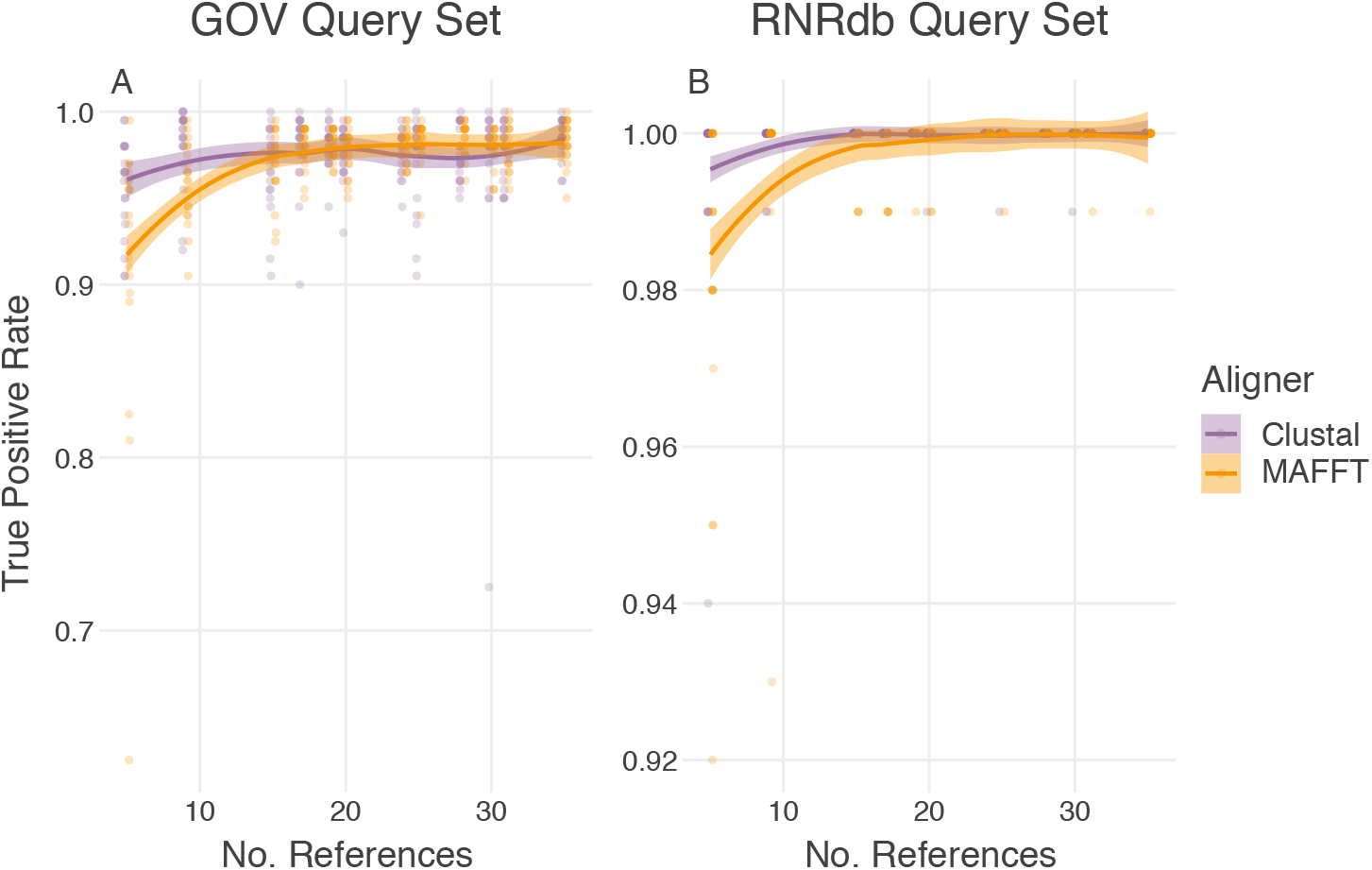
PASV true positive rate increases with number of references. Number of references per reference set versus PASV true positive rate for full-length reference sets. GOV query set and RNRdb query set are shown in panel A and panel B, respectively. Each dot represents a single PASV run (i.e., one reference set with an aligner: Clustal Omega – purple, MAFFT – orange). Locally estimated scatterplot smoothing (LOESS) lines with 95% confidence intervals are shown for each aligner. (Note the difference in y-axis scale between panels A and B.)

While using an increasing number of references boosted PASV accuracy, it also increased runtime (Additional File 1), as more sequences needed to be aligned. Using full-length references as opposed to references trimmed to the region of interest also increased the runtime. This is due to full-length references containing more bases that need to be aligned. Another consideration for run-time is the alignment algorithm: running PASV with Clustal Omega was faster than with MAFFT (Additional File 1).

In summary, variables that had the most impact on PASV true positive and true negative rate were alignment software (with Clustal Omega outperforming MAFFT) and reference trimming (full-length references performing better than references trimmed to the ROI) (Fig. 5, Table 1, Additional File 2).

### Testing PASV with the full GOV query set

PASV was tested on a large metagenomic query set using best practices determined from the 1,920 reference set tests. The only variable significantly associated with PASV accuracy was phylogenetic vs. random reference picking, which affected the true negative rate; however, the size difference was small (0.027%) (Additional File 3). As the different reference sets all had comparable results, the mean and 95% CI of all 18 reference set runs was used for the confusion matrix. Overall, PASV was highly concordant with the manual curation, with >99% agreement between PASV predictions and manual curation (Table 2). As in the full reference set tests, runtime increased with increasing numbers of reference sequences (Additional File 4) (linear model: runtime = (−13.5 *±* 6.8) + (5.26 *±* 0.3) * number of reference sequences).

**Table 2.** Confusion matrix of PASV results for 18 references sets against putative GOV RNR sequences.

| PASV Prediction | Manual curation |  |
| --- | --- | --- |
|  | Positive | Negative |
| Positive | 2894.6 ± 5.3 | 12.5 ± 0.8 |
| Negative | 21.4 ± 5.3 | 6977.5 ± 0.8 |

Mean ± 95% confidence interval for 18 PASV runs. Each run is one of 18 reference sets with the full 9,906 sequence Global Ocean Virome (GOV) query set.

PASV provides a means for automating the process of validating the identity of peptide sequences collected through homology search. The algorithm partitions query peptides into bonafide and by-catch sequences (Fig. 1). Given this, the impact of including by-catch sequences in a phylogenetic analysis of metagenomic RNR sequences was examined. Phylogenetic trees of putative RNR sequences from GOV (9,906 sequences), and sequences from the putative RNRs that PASV identified as bonafide (i.e., those sequences with N437, C439, E441, C462, *E. coli* numbering) were compared (Additional File 5). For this PASV run, the best performing reference set (hereby referred to as the "best practices" reference set) of the 18 tested on the full GOV query set that also followed the best practices observed in the full reference set test (i.e., full-length, single sequence per clade, random selection) was used. This PASV run yielded 2,914 bonafide RNR sequences (i.e., those sequences with N437, C439, E441, C462, *E. coli* numbering).

The tree including all putative RNRs contained a high proportion of sequences on long branches, indicative of distantly related sequences or sequences with poor alignment (Fig. 7A). In contrast, the bonafide PASV sequence tree contained fewer long branches and more reasonable topology [60] (Fig. 7B). In the case of both trees, clades with long branches did contain nontarget sequences such as helicases, DNA polymerases, terminase, and thioredoxin (Fig. 7 and Additional File 6). However, the tree containing bonafide RNR sequences had substantially fewer long branches, and those that were present would be relatively easy to identify and remove. In practice, having fewer long branches reduces the time necessary for manual curation of phylogenetic trees.

**Figure 7.**
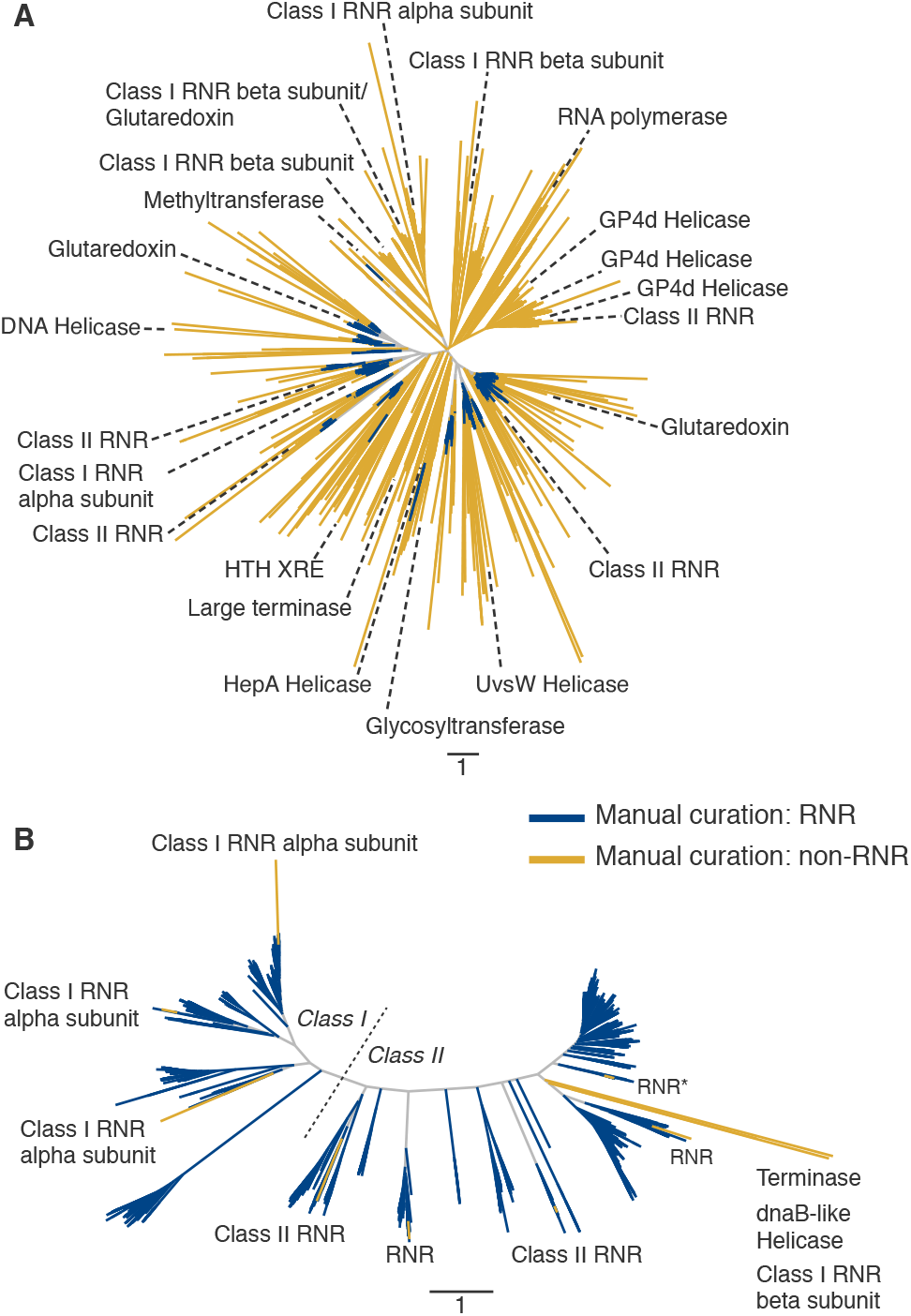
Phylogenetic trees of putative and bonafide GOV RNR sequences. Approximately-maximum likelihood trees of (A) 9,906 putative GOV RNR sequences identified by MMseqs2 using sensitive homology search parameters, and (B) 2,914 PASV validated, bonafide GOV RNR sequences (i.e., sequences with N437, C439, E441, C462, *E. coli* numbering). In panel B, the dotted line indicates the divide of Class I and Class II RNR sequences. Branch colors correspond to the results of manual curation. Blue branches indicate sequences manually annotated as RNR, whereas yellow branches represent sequences annotated as non-RNR or non-functional RNR sequences. Labelled sequences represent a sampling of sequences with homology to RNR, but manually curated as non-RNR or nonfunctional RNR. Note that some yellow branches in panel B, which were originally annotated as RNRs through manual curation, but having the correct residues according to PASV, were found to have correct RNR annotations according to the NCBI CDD [57]. The branch labeled "RNR*" in panel B indicates 3 branches annotated as RNR by the CDD.

Across all 18 GOV PASV runs (1 run per generated reference set), a total of 187 sequences out of 9,906 showed disagreement between PASV predictions and manual curation. These 187 sequences were annotated using NCBI CDD (Table 3, Additional File 3). Annotations of the 162 PASV predicted negative, manual curation positive sequences included three Class I RNR alpha subunits, 63 Class II RNRs, and 96 RNRs with unknown subclass. Sequences with hits to the RNR_PFL superfamily were considered to be either Class I alpha or Class II RNRs for two reasons: 1) other members of the supergroup, pyruvate formate lyase (PFL) and Class III RNRs, are oxygen-sensitive [61, 62], and thus unlikely to be found in the environments sampled in the GOV study [52]; and 2) these sequences grouped with other Class I alpha and Class II sequences on the phylogenetic trees (Fig. 7). The 25 remaining mismatched sequences (i.e., PASV predicted positive, manual curation negative) had more heterogeneous annotations. Twelve of these had hits to non-RNR domains: four helicases, three Pol Is, two endonucleases, one terminase, one Ankyrin repeat, and one with no match. Thirteen had hits to RNR domains: four Class I alpha subunit, two Class I beta subunit, two Class II RNR, and five RNRs with unknown subclass. The six sequences annotated as RNR Class I alpha subunits and Class II represent sequences that were likely erroneously categorized during manual curation. Thus, PASV was correct according to CDD annotations in 175 of the 187 cases in which PASV predictions disagreed with manual annotation.

**Table 3.** NCBI CDD annotations of sequences with mismatched PASV prediction and manual curation.

| Annotation | Count |  |
| --- | --- | --- |
|  | PASV positive, manual curation negative | PASV negative, manual curation positive |
| RNR (subclass unknown) | 5 | 96 |
| Class I RNR alpha subunit | *4 | 3 |
| Class I RNR beta subunit | 2 | - |
| Class II RNR | *2 | 63 |
| Helicase | 4 | - |
| Pol I | 3 | - |
| Endonuclease | 2 | - |
| Terminase | 1 | - |
| Ankyrin repeat | 1 | - |
| No match | 1 | - |

Counts are totals across 18 PASV runs: the full 9,906 sequence Global Ocean Virome query with 18 different reference sets.
\*Sequences erroneously categorized as non-RNR by manual curation

### Partitioning RNR Class I alpha subunit and Class II sequences

PASV’s ability to partition two biochemical classes of RNR sequences (Class I alpha subunit and Class II [63, 62]) was examined. The 2,579 RNRdb sequences used to make the RNR tree for phylogenetic clustering (Fig. 3) were partitioned into Class I alpha subunits and Class II sequences using PASV. As the NCEC residues within the RNR PASV profile are required for RNR function [45, 46, 47, 48], any sequence without NCEC at residues 437, 439, 441, and 462 (*E. coli* numbering) in the PASV run were grouped into the “Other” category. These included five Class I alpha and four Class II sequences. For the remaining 2,570 NCEC sequences, any sequence that PASV predicted as having an leucine at position 438 was labeled as a Class I alpha subunit, whereas any sequence with a proline at that position was predicted to be Class II. These PASV predictions were compared to RNRdb annotations, and the results were recorded in Table 4 (Additional File 7). Of the 1,244 annotated Class I alpha sequences, PASV predicted 1,236 of them to be Class I alpha (correct PASV predictions: 98.96%), one to be Class II (0.08%), and seven to be “Other” (0.96%). For the 1,326 annotated Class II sequences, PASV predicted 1,307 of them to be Class II (correct PASV predictions: 98.27%), four to be Class I alphas (0.30%), and 15 “Others” (1.43%).

**Table 4.** PASV Class I alpha and Class II predictions.

| PASV Prediction | Manual annotation |  |
| --- | --- | --- |
|  | Class I alpha | Class II |
| Class I alpha<sup>a</sup> | 98.96% | 0.30% |
| Class II<sup>b</sup> | 0.08% | 98.27% |
| Other<sup>c</sup> | 0.96% | 1.43% |

<sup>a</sup>NCEC sequences with L438 (*E. coli* numbering)
<sup>b</sup>NCEC sequences with P438 (*E. coli* numbering)
<sup>c</sup>Non-NCEC sequences or those with any other residue at position 438

### Partitioning AOX and PTOX sequences

A total of 336 alternative oxidase (AOX) and plastid terminal oxidase (PTOX) peptide sequences were recovered from a previous study examining misannotation of the AOX and PTOX gene groups in plants [37]. These sequences were classified with PASV using residues from the diagnostic, conserved motifs identified in [37]. This experiment tested the ability of PASV for correctly binning a mixed collection of AOX and PTOX peptide sequences. While distinct proteins, AOX and PTOX share regions of homology and are frequently missanotated by standard methods [37]. However, two motifs for each protein, when used in conjunction with MSA, enables correct classification of the proteins. Two reference sets were constructed, one to classify AOX and one to classify PTOX. The entire query set (336 total sequences, 254 AOX, 82 PTOX) was run through the PASV algorithm against both reference sets (Additional Files 8 & 9). In the AOX run, any sequence with the correct residues in the conserved motifs as identified by PASV was considered an AOX (motif 1: E233, R234, M235, H236, L237, M238, T239; motif 2: L283, E284, E285, E286, A287, numbered according to sequence O22048). Any sequence containing any other residue in any of these positions was considered to be non-AOX. In the PTOX run, sequences that PASV identified as having the correct residues in motifs 1 and 2 were annotated as PTOX (motif 1: G157, W158, R160, R161; motif 2: H177, H178, L179, L180, M182, E183, numbered according to sequence A0A061GHF5). Sequences containing different residues in any of these positions were annotated as non-PTOX. When these two annotations were combined, PASV correctly identified all 254 AOX and 82 PTOX peptide sequences and misannotated none.

## Discussion

Homology tools used for collecting gene sequences from databases and metagenomes, such as BLAST [21], HMMER [38], MMseqs2 [39], or PSI-BLAST [40], are sensitive and have the ability to detect remote homology between sequences. While detecting distant homologs is useful, especially when analyzing environmental metagenomic data, such sensitivity often comes with a price: increased levels of false positive sequences [27]. In the context of viral and microbial ecology, false positives can include non-functional versions of the protein of interest, correctly annotated proteins that do not span a predetermined region of interest, and proteins that share a conserved region or domain with the protein of interest, but are not the desired protein.

Including such false positives in analyses of functional proteins causes a number of problems. False positives interrupt multiple sequence alignments and subsequent phylogenetic analyses, which leads to inaccurate conclusions as to the evolutionary history of a protein [64, 65]. In ecological studies, inclusion of false positive sequences in marker gene phylogenetic analyses can lead to erroneous identification of microbial or viral populations [66, 37, 23].

Manual validation of proteins becomes increasingly error-prone and impractical with increasing dataset size. While larger datasets provide the means for deeper exploration of microbial communities and protein diversity and evolution, they also yield more protein sequences for validation. Sensitive homology searches can result in thousands of protein sequences from a single metagenome library, making automatic validation an attractive option.

### Using RNRs to test PASV

Any protein containing conserved residues, whether these are discovered purely through computational methods or are backed by biochemical characterization experiments can be validated using PASV. Ribonucleotide reductase (RNR), an ancient enzyme with well understood structural biochemical features [67] that is often misannotated in sequence databases [36], was an excellent experimental model for testing PASV’s ability to validate and partition putative RNR sequences collected from large sequence datasets by homology search. RNRs contain many immutable residues that have been discovered through decades of structural biology research [68]. There is at least one documented case of a gene with high sequence homology to RNR with mutated active sites that has evolved to perform an alternative function [69].

While RNRs are evolutionarily related, perform the same function, and are biochemically conserved, some share only 10-20% primary sequence similarity, a level below the “twilight zone” of homology search similarity [70, 71, 72]. Searching for RNRs, therefore, requires sensitive homology searches, which can return many false positive sequences. Due to the low level of sequence similarity among RNRs in general, and its many classes and subclasses, RNRs can be difficult to annotate. In one survey of RNRs recovered from GenBank, only 23% were deemed to be annotated correctly and 16% had not been annotated as RNRs at all [36]. Given the frequency of misannotation, low sequence homology, presence of immutable residues, and the RNRdb, a large, hand-curated database of bonafide RNR sequences [36], RNR provided an excellent model system for testing PASV. In addition, RNRs are of interest to researchers in many fields, including evolution, biochemistry, cancer research, and viral ecology [62, 18].

Here, we focused on Class I and II RNRs, which are the two most closely related extant RNRs. Class I RNRs are encoded by two genes, one each for the alpha and beta subunits comprising the active protein [67]. The larger alpha subunit is hypothesized to be the direct descendent of Class II RNRs [72], while the beta subunit belongs to the ferritin-like superfamily [73] and bears no homology to either Class I alpha or Class II RNRs. Class I and II RNRs require different cofactors for ribonucleotide reduction, so differentiating the classes is crucial for subsequent ecological analyses [18, 23].

PASV was tested using RNRs from two contrasting datasets: the RNRdb [36] and Global Ocean Viromes (GOV) [52]. The majority of RNRs in the RNRdb are from known organisms within large sequence databases (e.g. GenBank, SwissProt, etc.), with relatively few sequences originating from metagenomes. Virus sequences are relatively rare in curated databases as compared to sequences from eukaryotes and bacteria. In fact, viral sequences make up only 2.7% of the Class I alpha and Class II RNRs in the RNRdb. GOV, in contrast, is an environmental dataset of viral sequences. Thus the RNRdb and GOV represented different challenges for PASV.

### Factors influencing PASV accuracy

The most important factors influencing PASV accuracy surrounded the relative length of reference sequences and the approach used for choosing them. Using full length reference sequences, picking references randomly from a pool of potential sequences rather than based on phylogenies, and using more reference sequences all increased accuracy as measured by true positive and true negative rate. The benefit of using more reference sequences, however, plateaued after ca. 20 sequences in the reference set (Fig. 6), while the computing time required by PASV continued to increase (Additional File 1).

For each phylogenetically-informed reference set generated, a size-matched set of randomly selected RNRs were chosen to act as a control. It is important to note that while the randomly selected sequences are random with respect to their position on the tree, sequences from the RNRdb are biased with respect to class and subclass representation. Therefore, the “random” controls can also be seen as weighted by the composition of the RNRdb.

Alignment software was also a factor, with Clustal Omega generally outperforming MAFFT. However, this advantage was mostly lost when using full-length reference sequences rather than references trimmed to the region of interest. This result may also differ depending on the protein to be aligned, as some datasets are more difficult to align than others [74].

Reference sets representing as much of the known diversity of RNRs as possible (i.e., those taken evenly from across major clades of a phylogenetic tree) were hypothesized to increase PASV accuracy. This hypothesis was built on the idea that including diverse RNRs would prevent large irregularities in the alignments from more divergent query sequences. However, including diverse RNRs had the opposite effect and statistical tests showed that randomly selecting full-length reference sequences resulted in greater accuracy. One explanation for this phenomenon is that accuracy of multiple sequence alignment decreases with increasing sequence heterogeneity [35, 34]. As a consequence, forcing divergent sequences into the reference sets likely destabilized the alignments and decreased PASV’s accuracy.

### Using PASV to eliminate bycatch of non-target sequences

The GOV dataset provided an alternative experimental model for testing how PASV performed as a post-processing step after a homology search of a metagenomic sequence library. PASV effectively filtered out false-positive bycatch sequences recovered from the environmental metagenomes while searching for the gene of interest, RNR. Of the nearly 10,000 putative RNR sequences identified by MMseqs2, only about one-third were validated as functional RNRs by both PASV and manual curation. The other two-thirds were considered bycatch sequences. Common gene families within the bycatch sequences included RNR Class I beta subunits, thioredoxins, glutaredoxins, polymerases, helicases, and terminases (Additional File 6). Given the sensitivity of MMseqs2 [39], it is likely to find significant hits in sequences only distantly related to RNR or to sequences with domains similar to those occasionally found in RNRs. Some RNR Class I beta subunits are known to contain fused glutaredoxin domains [75]. RNRs may also have regions of remote homology to polymerases, helicases, and terminases as all of these proteins bind DNA. Some RNRs are known to contain zinc-finger domains [76], and at least one of the helicases examined with the CDD contained a zincfinger domain as well (Additional File 6).

Overall, PASV did an excellent job of removing most bycatch sequences (Table 2). Across the 18 reference set experiments that used the full GOV query set, only 187 of 9,906 RNR sequences had PASV predictions that disagreed with manual curation (Additional File 10). In most instances these sequences, annotated as terminases, polymerases, and helicases by NCBI CDD, existed on long branches indicating significant evolutionary distance from true Class I large subunit and Class II RNR sequences (Fig. 7B). Many of the false-positives identified by PASV (those sequences that PASV predicted to be RNRs, but manual curation predicted to be non-RNR) were likely RNR sequences that were missed during manual annotation. This can be attributed to the challenge of manually curating thousands of sequences and the problems inherent when performing large multiple sequence alignments.

### Partitioning sequences by key residues

PASV was conceived as a tool for validating the identity and functionality of protein sequences following homology searches. However, use cases for PASV extend beyond separation of bonafide and bycatch sequences. PASV provides an automated method for applying domain knowledge of a target protein to a large number of sequences. From this domain knowledge, PASV can partition sequences into groups based on structural characteristics that may be linked with protein biochemistry or phylogeny.

PASV was used in such a way to partition Class I alpha and Class II RNRs. While many amino acid residues in the active and allosteric sites of Class I alpha and Class II RNRs are conserved, other residues may be diagnostic of class [23]. Prior work based on protein alignments and phylogenetic trees suggests that the residue in position 438 (*E. coli* numbering) may be diagnostic of RNR class. Thus, we tested PASV’s ability to leverage this domain knowledge by sorting RNRdb sequences into class based on the identity of the residue in position 438. The function of residue 438 is unknown, but it is known to be conserved and sits within the active finger loop domain that contains the immutable active sites N437, C439, and E441 [77]. The sorting by PASV agreed almost perfectly with the RNRdb class annotations (Table 4), with >98% of Class I alpha and Class II sequences correctly identified.

An extension of this use case are peptides that cannot be differentiated by homology searches alone. Alternative oxidase (AOX) and plastid terminal oxidase (PTOX) are membrane-bound di-iron carboxylate proteins that oxidize a quinol substrate [78]. Although the proteins function within different organelles (AOX functions within the mitochondrial electron transport chain [79, 80] while PTOX is a chlororespiration enzyme only found within plastids and cyanobacteria [81]), their shared homology and function has led to high levels of misannotation [37]. However, using the amino acid signatures presented previously [37], PASV was able to sort AOX and PTOX proteins from each other with 100% accuracy. In this way, PASV leverages expert knowledge in an automated fashion.

We have shown that PASV can accurately partition Class I alpha and Class II RNRs using a residue diagnostic of these classes (Table 4), and AOX sequences from PTOX sequences using conserved motifs [37]. Given its success with these two disparate examples, it is likely that PASV could be effectively applied to other gene partitioning tasks as well. For example, a single amino acid mutation at position 762 (*E. coli* numbering) of motif B of DNA polymerase I (Pol I) imparts dramatic changes in either the fidelity or efficiency of replication [82]. Subsequent work has hypothesized that Pol I 762 mutations predict the life history characteristics [17] and the genetic composition of the replication module [14] of bacteriophages using Pol I for genome replication. PASV could be used to automatically partition viral Pol I sequences based on the 762 position, providing a means to further test hypothesized connections between Pol I biochemistry and phage life history using large metagenomic datasets. There are many examples of point mutation(s) in bacterial proteins that prevent antibiotics from binding and, thus, inhibit the function of the antibiotic (e.g., K88R in *rpsL* [83], C117D in *murA* [84], H526T in *rpoB* [85], Q124K in EF-Tu [86], V246A and V300G in *ndh* [87]). Such point mutations within a protein would not be readily apparent from homology search alone. Thus PASV could be used for validating and grouping these peptide sequences according to key point mutations following identification via homology search.

## Conclusions

Studies using gene sequences of functional proteins collected from metagenomes for investigating microbial diversity provide new challenges not faced when using genes for stable RNAs like SSU rRNA. These challenges include detecting and preventing false-positive bycatch sequences within datasets, validating key functional residues in proteins of interest, and partitioning peptide sequences into groups or classes. The PASV pipeline provides researchers with a means for addressing these challenges in an automated and highly accurate fashion by combining multiple sequence alignment with expert-curated domain knowledge. The PASV program and source code is freely available under the MIT license and can be found, along with documentation and usage examples, on GitHub: https://github.com/mooreryan/pasv.

## Availability of source code and requirements

- Project name: Protein Active Site Validation (PASV)
- Project home page: https://github.com/mooreryan/pasv
- Operating system(s): Any platform where Ruby and alignment software may be installed, or any platform that supports Docker
- Programming language: Ruby
- Other requirements: Ruby or Docker; alignment software, e.g., Clustal Omega, MAFFT, etc.
- License: MIT

## Availability of supporting data and materials

PASV source code and documentation are available on GitHub at https://github.com/mooreryan/pasv. The PASV Docker image is available on DockerHub at https://hub.docker.com/r/mooreryan/pasv. Data sets and miscellaneous scripts used in the preparation of the manuscript are available on Zenodo at https://doi.org/10.5281/zenodo.4426410. Additionally, a snapshot of the PASV source code v1.3.0 is available on Zenodo at https://doi.org/10.5281/zenodo.4426410.

## Declarations

## List of abbreviations

AOX: alternative oxidase
BRL: branch length
CDD: conserved domain database
GOV: global ocean virome
IQR: interquartile range
LOESS: locally estimated scatterplot smoothing
MSA: multiple sequence alignment
NCBI: National Center for Biotechnology Information
PASV: protein active site validation
PFL: pyruvate formate lyase
Pol I: DNA polymerase I
PTOX: plastid terminal oxidase
RNR: ribonucleotide reductase
ROI: region of interest

## Ethical Approval

Not applicable.

## Consent for publication

Not applicable.

## Author’s Contributions

- RMM: Conceptualization, Data curation, Formal analysis, Software, Writing – original draft, Writing – review & editing
- AOH: Conceptualization, Data curation, Writing – review & editing
- DJN: Conceptualization, Writing – review & editing
- JC: Conceptualization, Writing – review & editing
- MC: Conceptualization, Data curation, Visualization
- BDF: Conceptualization, Supervision, Writing – review & editing
- SWP: Conceptualization, Funding acquisition, Supervision, Writing – review & editing
- KEW: Conceptualization, Funding acquisition, Supervision, Writing – review & editing

## Acknowledgements

We would like to acknowledge current and past members of the Viral Ecology and Informatics Lab at the University of Delaware who tested early versions of PASV and provided valuable feedback: Jacob T Dums (ORCID: 0000-0002-6314-4779), Zach Schreiber (ORCID: 0000-0002-6271-2754), and Michael Dahle (ORCID: 0000-0003-0518-3355).

