## Additional File 1 for "PASV: Automatic protein partitioning and validation using conserved residues"

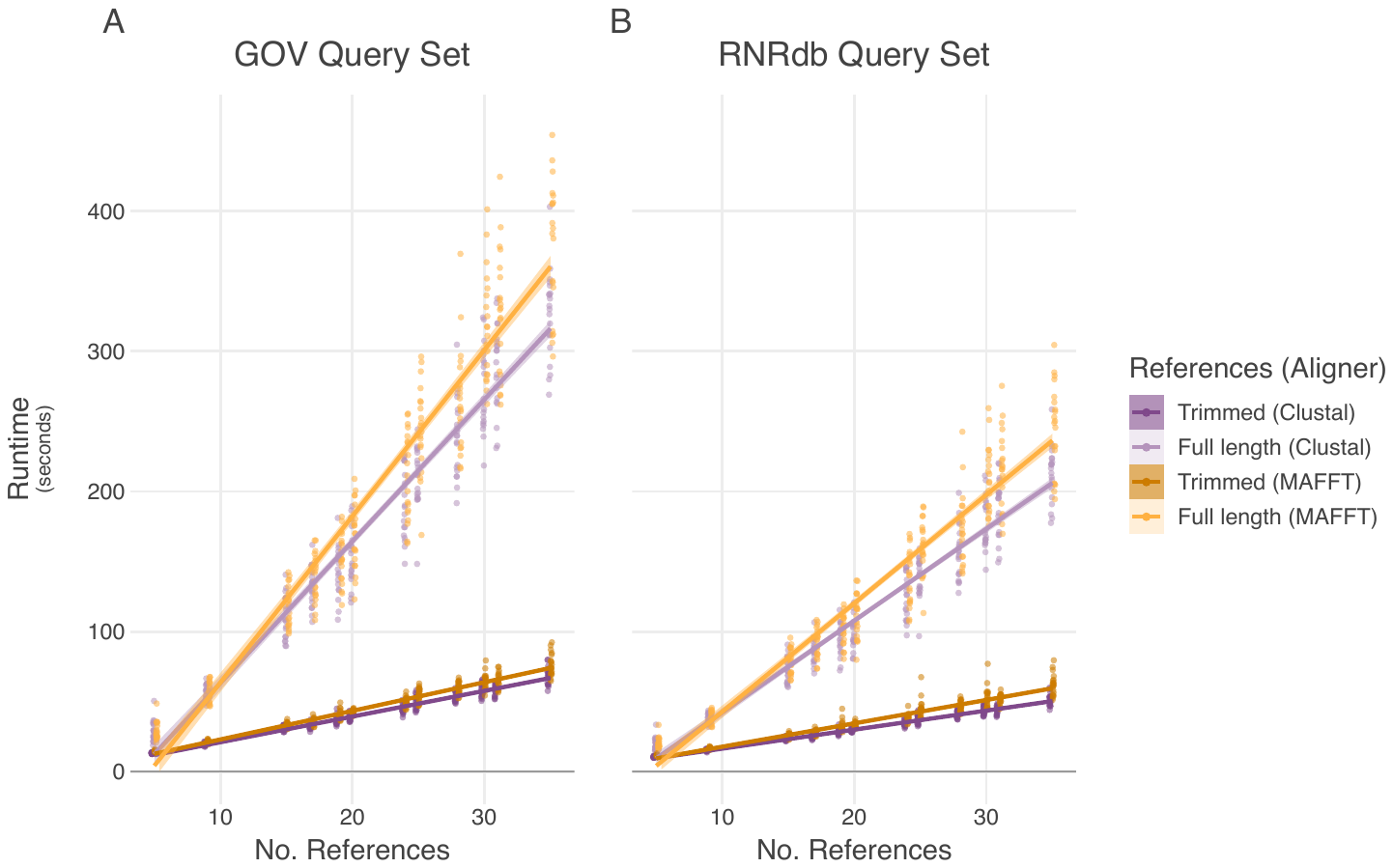

**Additional File 1: PASV runtime scales with number of references, reference length, and aligner.**

PASV runtime (walltime, in seconds) across 1,920 experiments.  Runtimes for the GOV (panel A) and RNRdb (panel B) query sets are shown separately as they contain a different number of queries (300 for GOV, 200 for RNRdb).  Each dot represents a single PASV run.  Dots are colored by reference length--aligner combinations (trimmed references -- light, full length references -- dark; Clustal Omega -- purple, MAFFT -- orange) and jittered for increased visibility.  Linear regression line with 95% confidence interval is shown.  All experiments were run on an Intel(R) Xeon(R) CPU E5-2695 v4 @ 2.10GHz server with 36 cores (2 threads per core) and 512 GB of ram, with PASV set to use 68 threads (i.e., process 68 queries concurrently).
