## Additional File 3 for "PASV: Automatic protein partitioning and validation using conserved residues"

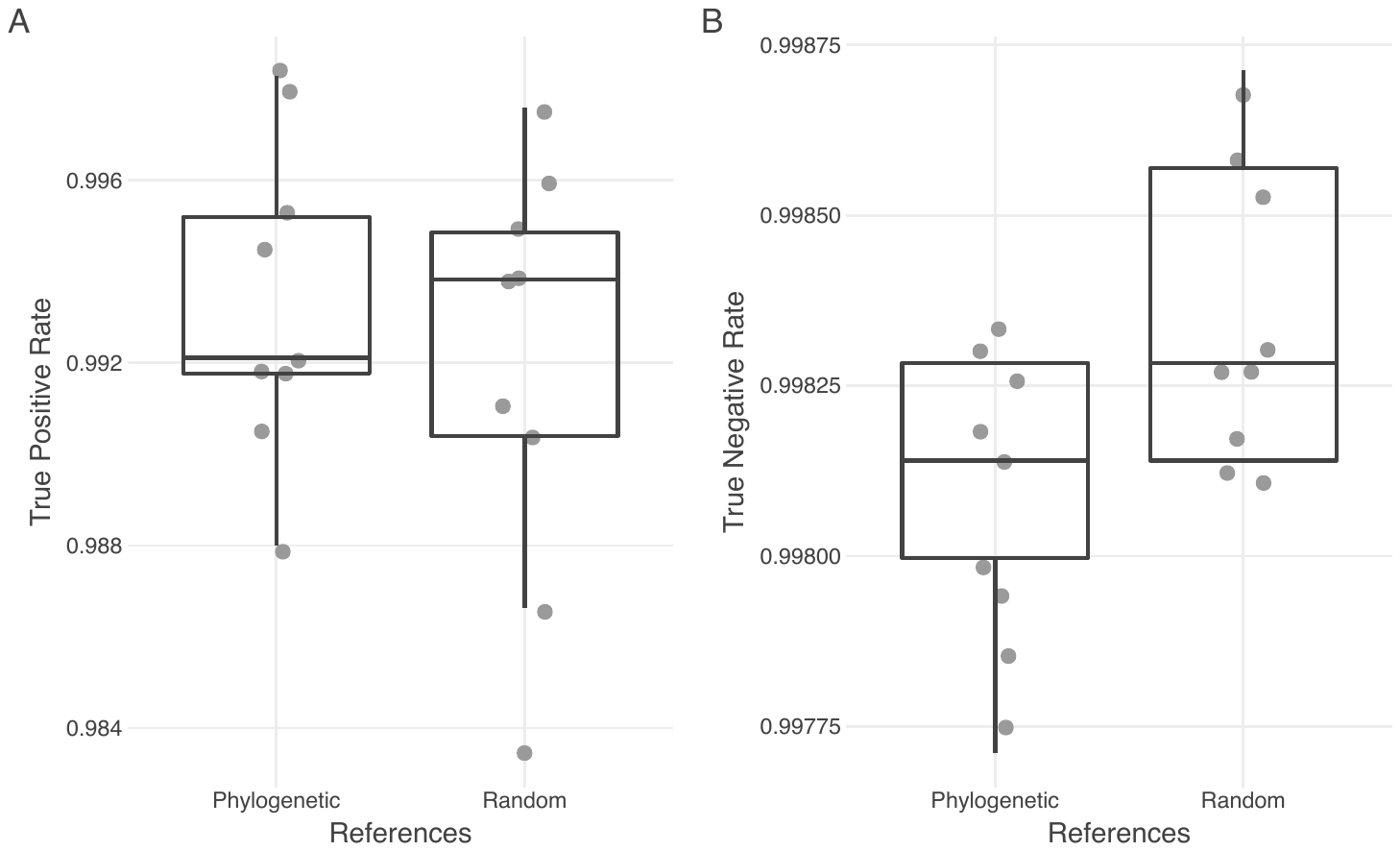

**Additional File 3: Phylogenetic vs. random selection of references on PASV performance for putative GOV RNR sequences.**

PASV performance (true positive rate -- panel A; true negative rate -- panel B) using Clustal Omega for eight reference sets against 9,906 putative GOV RNR sequences.  Dots represent individual PASV runs.  Box (showing median and interquartile range (IQR)) and whisker (showing 1.5 x IQR) plots are overlaid.  The difference between phylogenetic and random references is significant (according to a linear model with replicate, number of references, and phylogenetic vs. random reference selection; E-value < 0.05), though weak, for true negative rate.  (Note the difference in y-axis scale between panels A and B.)
