## Additional File 4 for "PASV: Automatic protein partitioning and validation using conserved residues"

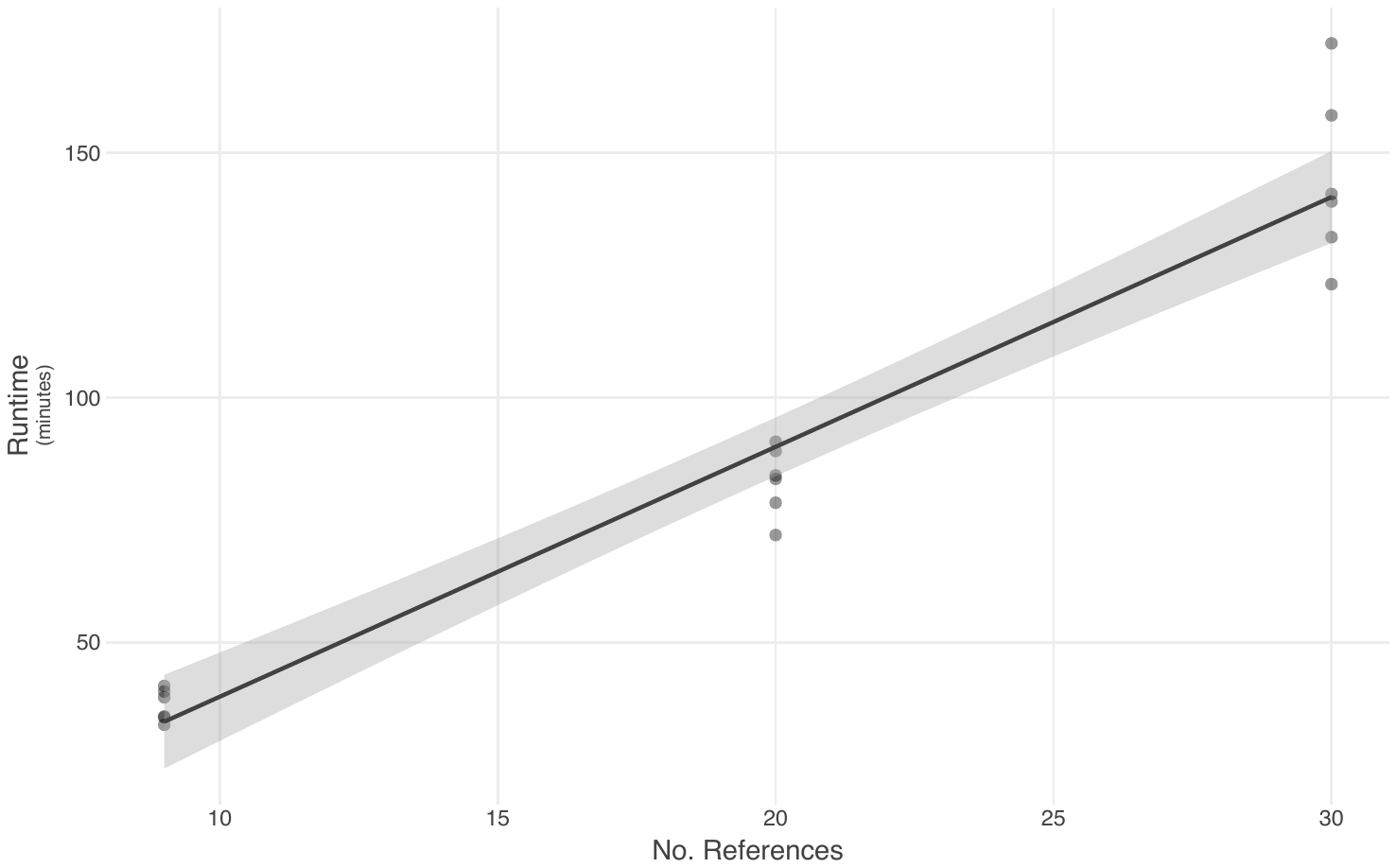

**Additional File 4: PASV runtime for putative GOV RNR sequences scales with number of references.**

PASV runtime (walltime, in minutes) for the full GOV query set (9,906 putative RNR sequences) across 18 experiments.  Each dot represents a single PASV run.   Linear regression line with 95% confidence interval is shown.  All experiments were run on an Intel(R) Xeon(R) CPU E5-2695 v4 @ 2.10GHz server with 36 cores (2 threads per core), with PASV set to use 68 threads (i.e., process 68 queries concurrently).
