## Additional File 5 for "PASV: Automatic protein partitioning and validation using conserved residues"

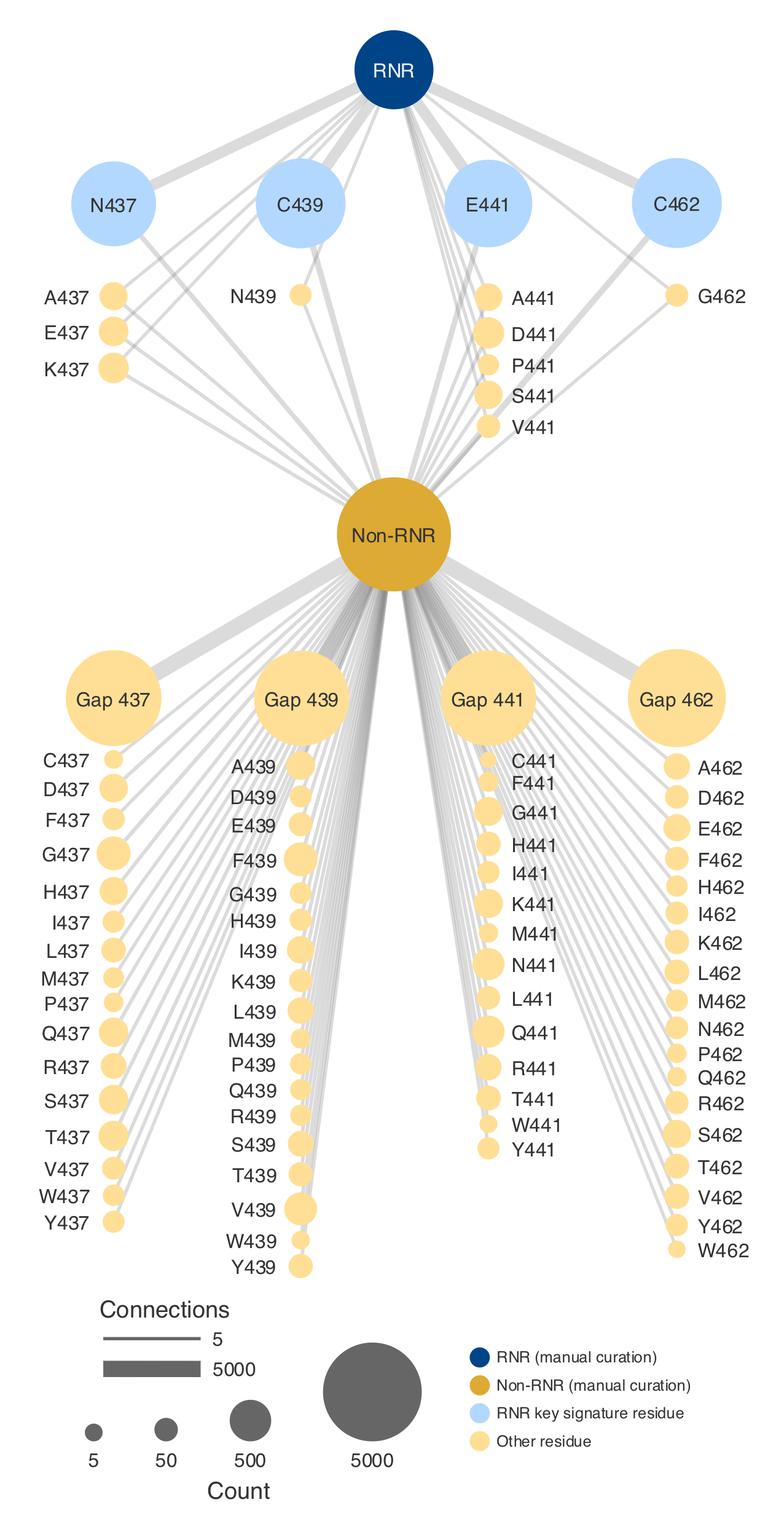

**Additional File 5: Network diagram of Global Ocean Virome RNR vs. non-RNR experiment.**

Putative ribonucleotide reductase (RNR) sequences from the Global Ocean Virome (GOV) (9,906 sequences) were manually curated into bins of RNR and bycatch. These sequences were then run through PASV using key signature positions 437, 439, 441, and 462. Results of the PASV run are visualized in the network. The group nodes (dark blue: manually curated RNR, and dark yellow: manually curated non-RNR) represent the result of the manual curation. The remaining nodes (i.e., residue-position nodes) represent amino acid residues (or a gap) found at a particular position in one of the key residue columns. Blue residue-position nodes indicate residue-positions included in the key signature (N437, C439, E441, and C462), and yellow nodes show other residue-position combinations found in the data. Node size for group nodes represent number of sequences in each of the two groups. Node size for the residue-position nodes represent the amount of times that residue-position combination was seen in the PASV output (e.g., N437 is larger than A437, indicating that N437 was more common in the data than A437). Residue-position nodes are linked to group nodes if that residue-position combination was seen in the corresponding group, with link thickness indicating the number of connections. For example, N437 (top-left corner) was seen in both RNR and non-RNR sequences (it is linked to both nodes), but more often connected with RNR sequences than with non-RNR sequences (the link between N437 and RNR is thicker than the link between N437 and non-RNR).
