## Additional File 7 for "PASV: Automatic protein partitioning and validation using conserved residues"

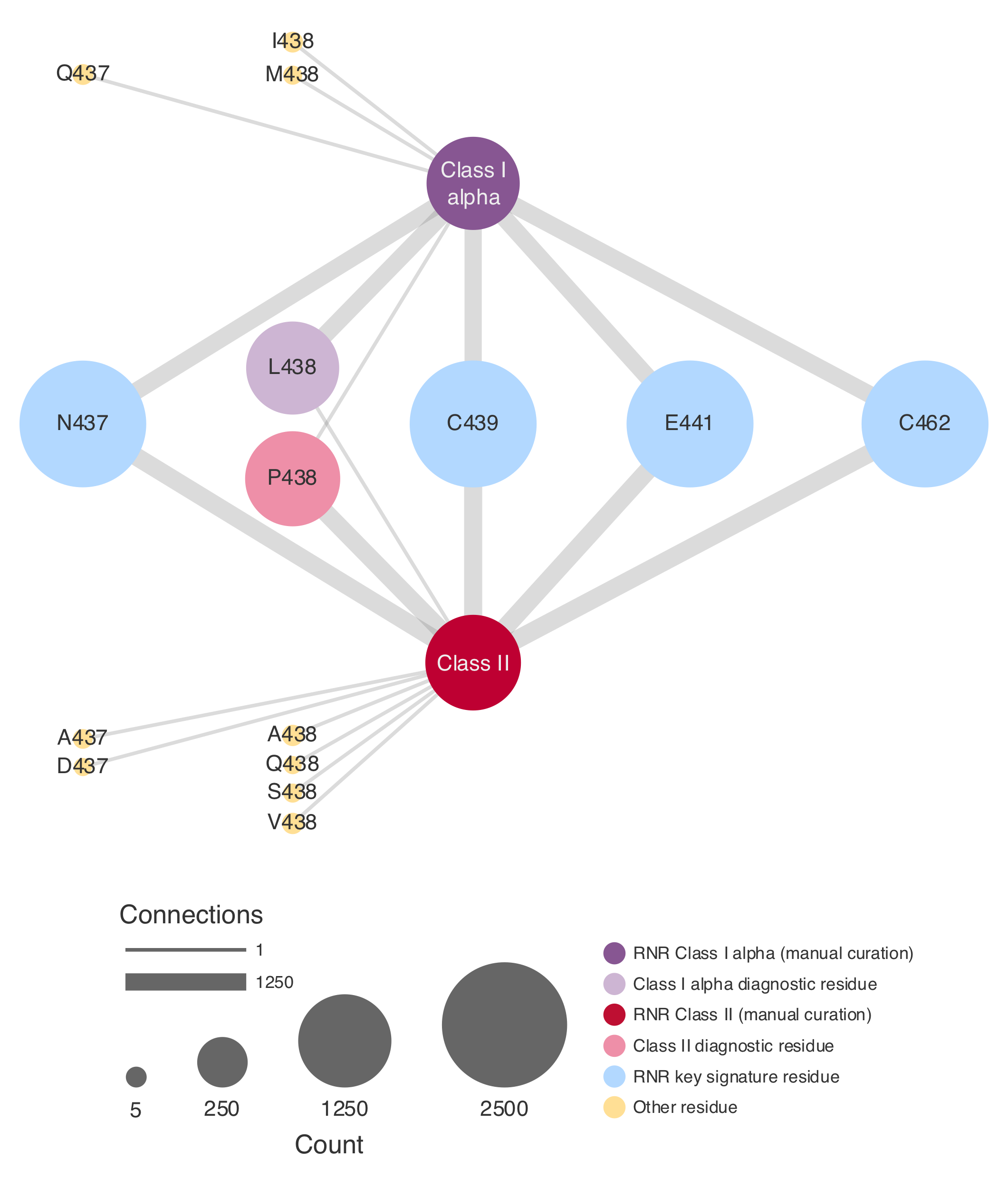

**Additional File 7: Network diagram of RNRdb Class I alpha vs. Class II RNR experiment.**

Class I alpha and Class II ribonucleotide reductase (RNR) RNRdb (2,579 sequences) were run through PASV using key signature positions 437, 438, 439, 441, and 462. Results of the PASV run are visualized in the network. The group nodes (purple: manually curated Class I alpha, and red: manually curated Class II) represent the class annotation of sequences included in the experiment. The remaining nodes (i.e., residue-position nodes) represent amino acid residues (or a gap) found at a particular position in one of the key residue columns. Blue residue-position nodes indicate required RNR residue-positions included in the key signature (N437, C439, E441, and C462), and yellow nodes show other residue-position combinations found in the data. The light purple node (L438) is the Class I alpha diagnostic residue-position, and the light red node (P438) is the Class II diagnostic residue-position. Node size for group nodes represent number of sequences in each of the two groups. Node size for the residue-position nodes represent the amount of times that residue-position combination was seen in the PASV output (e.g., N437 is larger than A437, indicating that N437 was more common in the data than A437). Residue-position nodes are linked to group nodes if that residue-position combination was seen in the corresponding group, with link thickness indicating the number of connections. For example, L438 (center-left) was seen in both Class I alpha and Class II RNR sequences (it is linked to both nodes), but more often connected with Class I alpha sequences than with Class II sequences (the link between L438 and Class I alpha is thicker than the link between L438 and Class II).
