## Additional File 8 for "PASV: Automatic protein partitioning and validation using conserved residues"

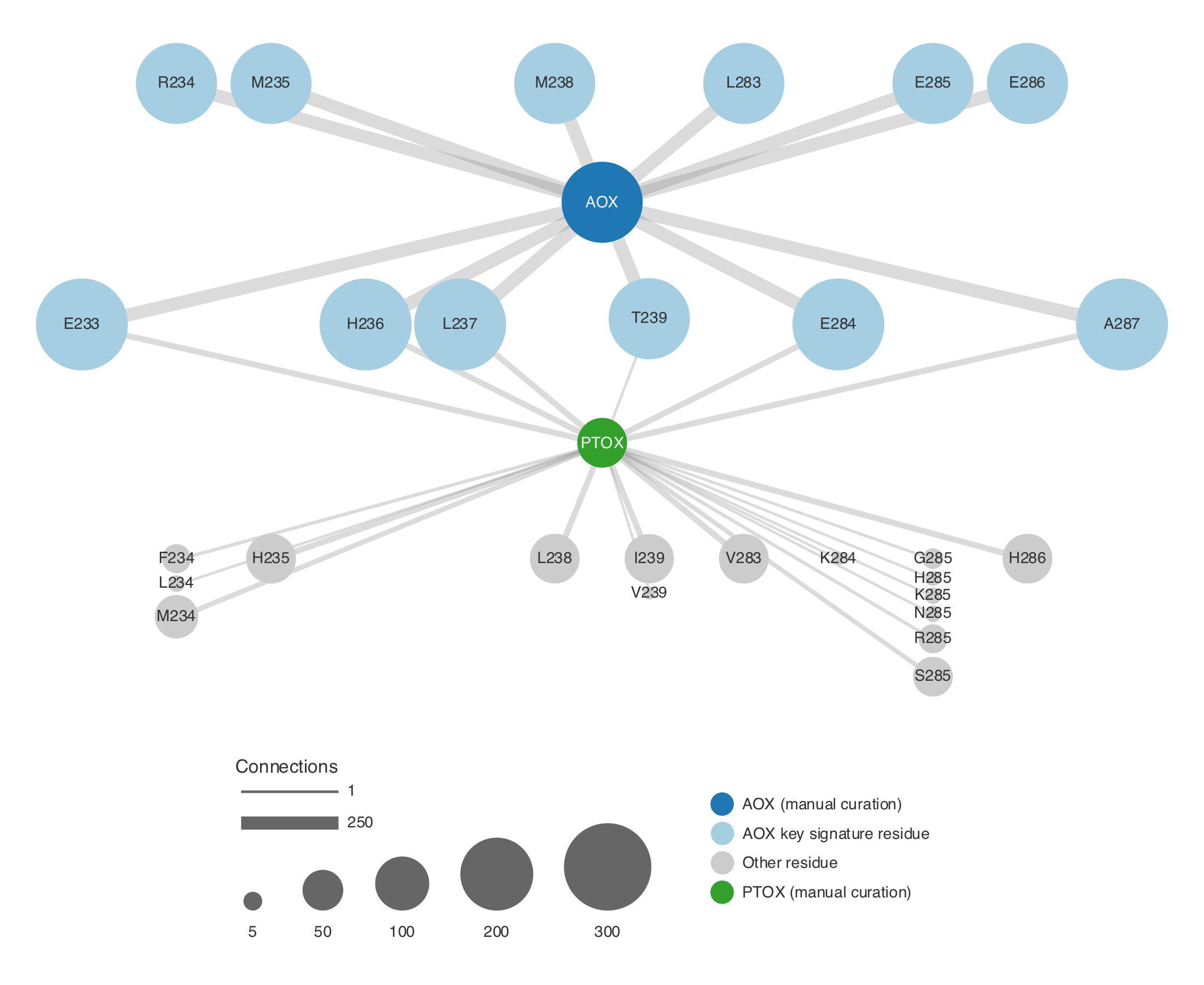

**Additional File 8: Network diagram of AOX-PTOX experiment using the AOX key signature and references.**

Alternative oxidase (AOX) and plastid terminal oxidase (PTOX) peptide sequences (336 total) were run through PASV using the AOX reference set and key signature positions 233, 234, 235, 236, 237, 238, 239, 283, 284, 285, 286, 287. Results of the PASV run are visualized in the network. The group nodes (blue: manually curated AOX, and green: manually curated PTOX) represent the annotation of sequences included in the experiment. The remaining nodes (i.e., residue-position nodes) represent amino acid residues (or a gap) found at a particular position in one of the key residue columns. Light blue residue-position nodes indicate correct residue-position combinations from AOX motifs 1 (E233, R234, M235, H236, L237, M238, T239) and 2 (L283, E284, E285, E286, A287), and gray nodes show other residue-position combinations found in the data. Node size for group nodes represent number of sequences in each of the two groups. Node size for the residue-position nodes represent the amount of times that residue-position combination was seen in the PASV output (e.g., T239 (center) is larger than V239, indicating that T239 was more common in the data than V239). Residue-position nodes are linked to group nodes if that residue-position combination was seen in the corresponding group, with link thickness indicating the number of connections. For example, T239 (centr) was seen in both AOX and PTOX sequences (it is linked to both nodes), but more often connected with AOX sequences than with PTOX sequences (the link between T239 and AOX is thicker than the link between T239 and PTOX).
