## Additional File 9 for "PASV: Automatic protein partitioning and validation using conserved residues"

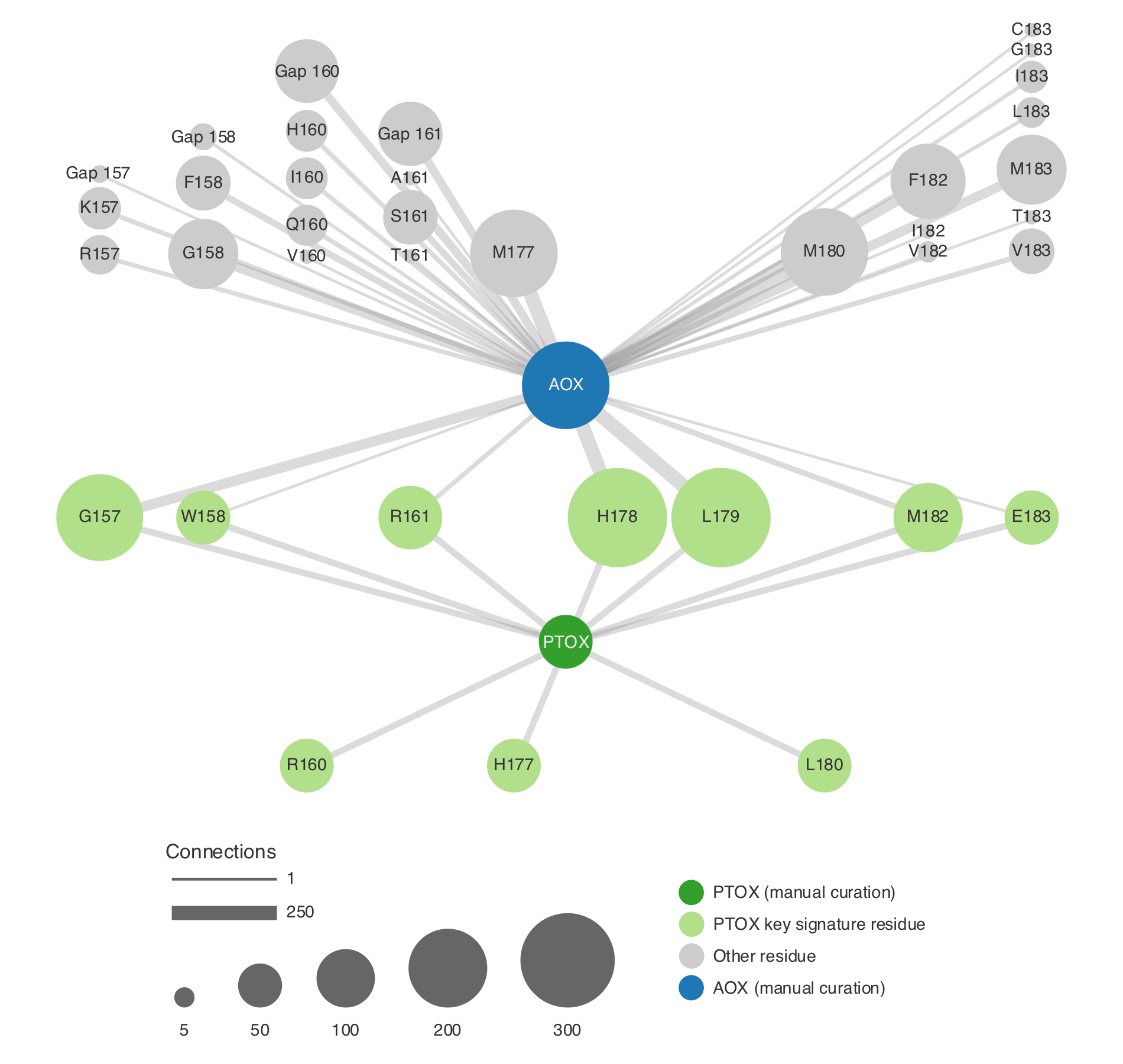

**Additional File 9: Network diagram of AOX-PTOX experiment using the PTOX key signature and references.**

Alternative oxidase (AOX) and plastid terminal oxidase (PTOX) peptide sequences (336 total) were run through PASV using the PTOX reference set and key signature positions 157, 158, 160, 161, 177, 178, 179, 180, 182, 183. Results of the PASV run are visualized in the network. The group nodes (blue: manually curated AOX, and green: manually curated PTOX) represent the annotation of sequences included in the experiment. The remaining nodes (i.e., residue-position nodes) represent amino acid residues (or a gap) found at a particular position in one of the key residue columns. Light green residue-position nodes indicate correct residue-position combinations from PTOX motifs 1 (G157, W158, R160, R161) and 2 (H177, H178, L179, L180, M182, E183), and gray nodes show other residue-position combinations found in the data. Node size for group nodes represent number of sequences in each of the two groups. Node size for the residue-position nodes represent the amount of times that residue-position combination was seen in the PASV output (e.g., E183 (middle-right) is larger than T183, indicating that E183 was more common in the data than T183). Residue-position nodes are linked to group nodes if that residue-position combination was seen in the corresponding group, with link thickness indicating the number of connections. For example, E183 (middle-right) was seen in both AOX and PTOX sequences (it is linked to both nodes), but more often connected with PTOX sequences than with AOX sequences (the link between E183 and PTOX is thicker than the link between E183 and AOX).
